# Alligamycin A, an unprecedented antifungal β-lactone spiroketal macrolide from *Streptomyces iranensis*

**DOI:** 10.1101/2024.04.17.589928

**Authors:** Zhijie Yang, Yijun Qiao, Emil Strøbech, Jens Preben Morth, Grit Walther, Tue Sparholt Jørgensen, Kah Yean Lum, Gundela Peschel, Miriam A. Rosenbaum, Viola Previtali, Mads Hartvig Clausen, Marie Vestergaard Lukassen, Charlotte Held Gotfredsen, Oliver Kurzai, Tilmann Weber, Ling Ding

**Author notes:** Corresponding author: Ling Ding. Contribute equally (Zhijie Yang, Yijun Qiao).

## Abstract

Fungal infections pose a great threat to public health and there are limited antifungal medicaments. *Streptomyces* is an important source of antibiotics, represented by the clinical drug amphotericin B. The rapamycin-producer *Streptomyces iranensis* harbors an unparalleled Type I polyketide synthase, which codes for a novel antifungal macrolide alligamycin A (**1**), the structure of which was confirmed by NMR, MS, and X-ray crystallography. Alligamycin A harbors an undescribed carbon skeleton with 13 chiral centers, featuring a (β-lactone moiety, a [6,6]-spiroketal ring, and an unprecedented 7-oxo-octylmalonyl-CoA extender unit incorporated by a potential novel crotonyl-CoA carboxylase/reductase. The *ali* biosynthetic gene cluster was confirmed through CRISPR-based gene editing. Alligamycin A displayed profound antifungal effects against numerous clinically relevant filamentous fungi, including *Talaromyces* and *Aspergillus* species. (β-Lactone ring is essential for the antifungal activity and alligamycin B (**2**) with disruption in the ring abolished the antifungal effect. Proteomics analysis revealed alligamycin A potentially disrupted the integrity of fungal cell walls and induced the expression of stress-response proteins in *Aspergillus niger*. Alligamycins represent a new class of potential drug candidate to combat fungal infections.

## Introduction

About 1.2 billion people worldwide are estimated to suffer from a fungal infection.^1^ Invasive fungal diseases are prevalent among immunocompromised populations, such as patients with HIV/AIDS, chronic lung diseases, prior tuberculosis, cancer, diabetes, and patients receiving immunosuppressant treatment and invasive medical procedures.^2^ In particular, invasive fungal diseases are associated with unacceptably high mortality rates. A recent report suggested an annual incidence of 6.5 million invasive fungal infections and 3.8 million deaths.^3^ For example, infections with *Aspergillus* have emerged as one of the most common death causes in severely immunocompromised patients, i.e. with acute leukemia and recipients of hematopoietic stem cell transplantation with mortality rates up to 40% to 50%.^1^ Furthermore, new risk cohorts are emerging, including patients with severe respiratory virus infection such as influenza or COVID-19.^4^ Currently, four classes of systemic antifungal medicines (azoles, echinocandins, pyrimidines, and polyenes) are used in clinical practice, and only a limited number of candidates are in the clinical development pipeline.^5–8^ While existing antifungal medications are effective in numerous instances, challenges persist due to side effects and complications arising from drug-drug interactions.^9^ Moreover, the emergence of resistance had rendered some existing medications ineffective, and this was partly driven by the inappropriate use of antifungals.^10^ For example, agricultural use of azoles has been linked to rising rates of azole-resistant *Aspergillus fumigatus* infections, with resistance rates of 15-20% reported in parts of Europe and exceeding 80% in environmental samples from Asia.^10–12^

*Streptomyces* are producers of bioactive small molecules exhibiting a broad range of structural and functional diversity called secondary or specialised metabolites. Secondary metabolites (SMs) are of great relevance to human health, offering pharmaceutical properties such as antibacterial, antifungal, anticancer, and immunosuppressive activities. Remarkably, in all new drug approvals from 1981 to 2019, over 60% of the 1394 small molecule drugs were either secondary metabolites or derivatives thereof.^13–15^ Nearly two-thirds of antibiotics approved for clinical use originate from *Streptomyces*, and polyketides represent a large and diverse group of *Streptomyces*-derived SMs. Many possess valuable antifungal properties, among which nystatin and amphotericin B are two active polyenes used clinically as first-line drugs to treat fungal infections. The first example is the antifungal drug nystatin isolated from *Streptomyces noursei* in 1950 by Hazen and Brown.^16^ Amphotericin B, isolated from *Streptomyces nodosus* in 1955, remains a crucial component in the therapeutic arsenal for combating invasive fungal diseases and leishmaniasis.^17^ Recent studies revealed that the mode of action of polyenes antibiotics involves acting as sterol “sponges” and extracting ergosterol from the fungal membrane, causing cell death, in addition to the classical model of pore formation in the fungal cell membrane.^18–20^ Amphotericin B has been used to treat mucormycosis, aspergillosis, blastomycosis, candidiasis, coccidioidomycosis, and cryptococcosis. However, adverse effects from amphotericin B treatment are common, with nephrotoxicity being the most serious.^21,22^ There has been a continued interest in finding new antifungal metabolites, such as turonicin A, hygrobafilomycin, iseolides, cyphomycin, azalomycins, niphimycins, and resistomycin.^23–29^ However, most exhibit non-selectivity and display toxicity to human cells. Thus, it is of clinical importance to develop effective selective antifungal drug candidates, with a new mode of action and low cytotoxicity.

Given the significant number of biosynthetic gene clusters (BGCs) present in *Streptomyces*, a plethora of their metabolites remains undiscovered.^30^ Access to modern analytical chemistry techniques in combination with genetics and bioinformatics tools has enabled us to revisit the “talented” microbes discovered decades ago.^31^ *Streptomyces iranensis*, initially identified as a rapamycin producer, was originally isolated from soil in Isfahan City, Iran.^32^ Subsequent investigations revealed its great capability to produce elaiophylin, azalomycins, nigericin, and other new metabolites yet to be discovered.^33^ In our recent research, we discovered that *S. iranensis* produced pteridic acids H and F, which could alleviate abiotic stresses efficiently during plant growth.^33^ During our pipeline search for antifungal polyketides, *S. iranensis* exhibited profound antifungal activity in co-cultivation with *Aspergillus flavus*, *Aspergillus niger*, *Aspergillus fumigatus and Aspergillus tubingensis* (*Supplementary information* Fig. 1), suggesting the production of antifungal metabolites. LC-MS analysis revealed a high production of the well-known antifungal azalomycin. However, the azalomycin-deficient mutant of *S. iranensis* still exhibited strong antifungal activity against *Aspergillus flavus*, *Aspergillus niger*, *Aspergillus fumigatus and Aspergillus tubingensis* (*Supplementary information* Fig. 1), which indicated the production of other potential novel antifungal metabolites. Genome mining of *S. iranensis* revealed the presence of other uncharacterized BGCs including a type-I modular polyketide BGC (*ali*) featuring sixteen modules. The product of this BGC namely alligamycin was identified by metabolomic analysis, large-scale fermentation and isolation. The structure and absolute configuration of alligamycins were elucidated using nuclear magnetic resonance (NMR), mass spectrometry (MS), and X-ray crystallography. The striking structural features of alligamycin A include a [6,6]-spiroketal ring, a rare β-lactone moiety, and a distinctive aliphatic branch derived from an unusual polyketide synthetase (PKS) extender unit. Alligamycin A exhibited potent antifungal activity, potentially due to disrupting fungal cell wall integrity, as suggested by label-free quantitative proteomics analysis. The β-lactone moiety appeared to be essential for the antifungal effect as alligamycin B featuring a disrupted β-lactone ring lost the antifungal activities. This study provides a paradigm for mining novel molecules from extensive natural microorganisms, with the potential to lead to the discovery of innovative antifungal agents to combat antimicrobial resistance.

## Results and Discussions

### Genome mining in *S. iranensis*

To obtain complete genomic data for *S. iranensis*, we conducted whole-genome resequencing by integrating the Oxford Nanopore Technologies MinION and Illumina MiSeq system. The high-resolution, full-genome sequencing of *S. iranensis* reveals a linear chromosome spanning 12,213,033 nucleotides, featuring inverted terminal repeats comprising 156,145 nucleotides. The BGC annotation of the acquired *S. iranensis* genome was carried out using antiSMASH version 7.0,^34^ which resulted in the identification of a cryptic BGC (*ali*) situated at the terminal region of the linear chromosome (Fig. 1a and *Supplementary information* Fig. 2). The gene cluster family (GCF) annotation of *ali* gene cluster showed it belongs to GCF_00315 and a total of 11 hits were detected (distance ≤ 900.0) in the BiG-FAM database (*Supplementary information* Tab. 2).^35^ The distinctive architecture of the biosynthetic genes in the *ali* BGC suggests its novelty as a modular Type-I PKS BGC, setting it apart from others (*Supplementary information* Fig. 3). Furthermore, the *ali* BGC demonstrated its uniqueness when compared with other BGCs in the NPDC database (https://npdc.rc.ufl.edu) as well as in house database.

**Fig. 1.**
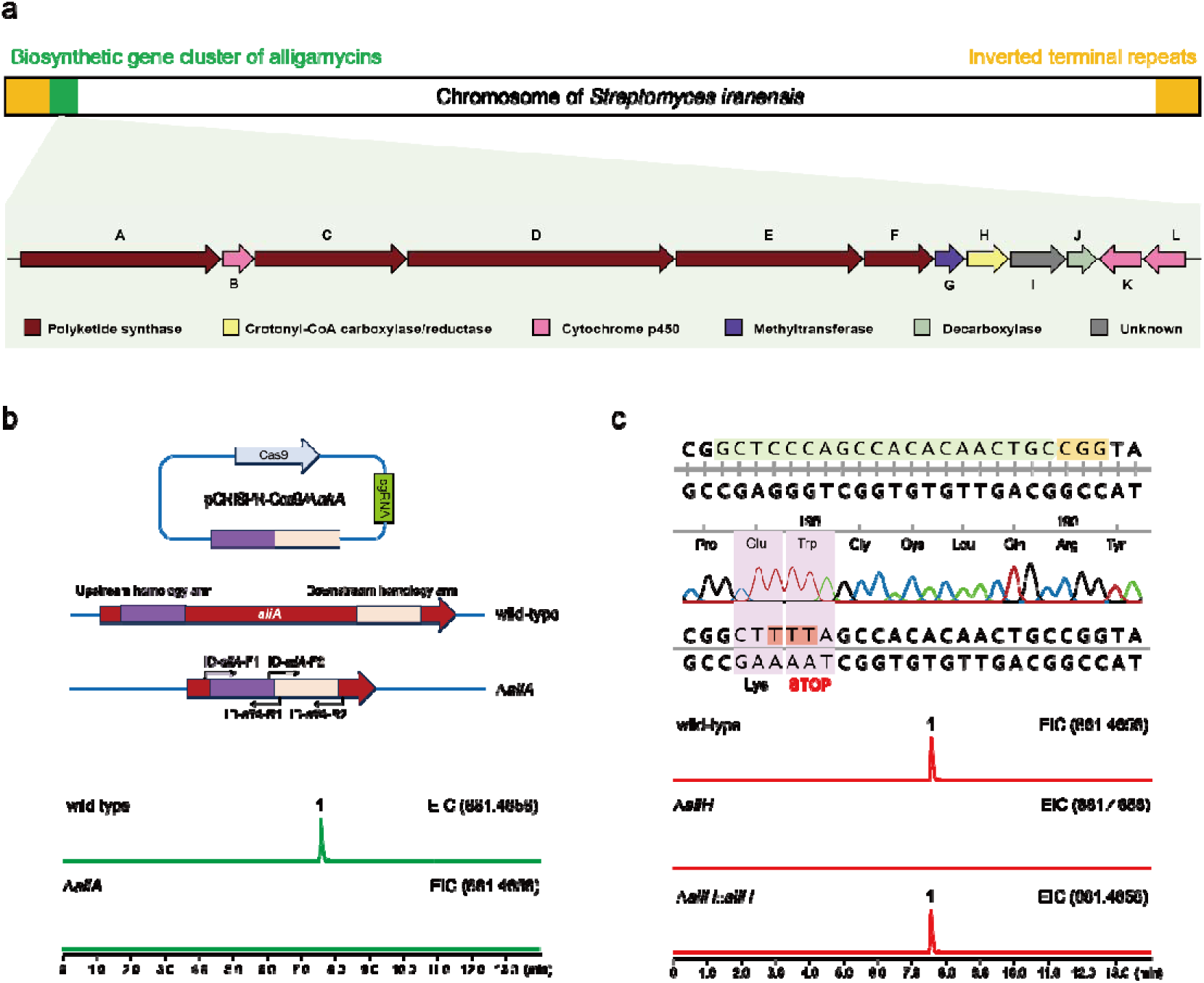
Organization of the alligamycin biosynthetic gene cluster (*ali*) and genome editing in *S. iranensis*. a) The *ali* BGC is situated at the end of the linear chromosome, adjacent to the inverted terminal repeats. Open reading frames are shown as arrows indicating the size and direction of transcription. b) Confirmation of the *ali* BGC through knock-out of *aliA.* Extracted ion chromatogram (*m/z* 881.4656 [M + Na]^+^) showing the abolishment of production of alligamycin in the Δ*aliA* mutant compared to the wild type strain. c) Inactivation of *aliH* in *S. iranensis* via CRISPR base editing. Extracted ion chromatogram (*m/z* 881.4656 [M + Na]^+^) showing the abolishment of production of alligamycin in the mutant *ΔaliH* compared to the wild strain. The production of alligamycin was resumed in complementation mutant *ΔaliH::aliH*.

To identify the potential products of the *ali* BGC, we combined both genome editing with metabolomics approaches. Initially, we constructed PKS knock-out mutant (Δ*aliA*) and crotonyl-CoA carboxylase/reductase-inactivated mutant (Δ*aliH*) using the CRISPR-Cas9 method and CRISPR base editing,^36^ respectively (Fig. 1). Comparative metabolic profiling using LC-MS and LC-UV led to the identification of compound **1** with *m/z* 881.4656 ([M + Na]^+^), which was absent in both mutants (Fig. 1b, 1c). The complementation of pGM1190-based *aliH* in mutant *S. iranensis*/Δ*aliH* led to the restoration of the production of **1** (Fig. 1c).

### Structure elucidation of alligamycins

To obtain the products of the *ali* BGC, a 200 L fermentation was carried out followed by a downstream processing using different chromatography techniques, including Amberchrom 161c resin, silica gel, Sephadex LH-20, and high-performance liquid chromatography (HPLC) to yield pure alligamycin A (**1**) and alligamycin B (**2**). The structures were elucidated using HR-ESI-MS, NMR spectroscopy, and X-ray crystallography.

Alligamycin A (**1**) was obtained as a white solid with a molecular formula of C_47_H_70_O_14_ as determined by HR-ESI-MS data (calcd *m/z* 881.4658 [M + Na]^+^). The ^1^H NMR spectrum revealed signals for six methyl groups Me-39 (δ 2.15), Me-41 (δ 0.94), Me-43 (δ 0.98), Me-44 (δ 0.93), Me-45 (δ 2.10), and OMe-8 (δ 3.37). Additionally, four olefinic protons were observed for H-2 (δ 6.19), H-3 (δ 7.91), H-4 (δ 6.47) and H-28 (δ 6.48). The coupling constant between H-2 and H-3 (*J =* 15.4 Hz) confirmed a trans-orientation of the double bond. Besides the presence of several oxygen-bearing methines (H-6, H-8, H-10, H-14, H-16, and H-22), various aliphatic proton signals were also observed (*Supplementary information* Fig. 6).

The ^13^C NMR spectrum showed signals for several carbonyl groups, one carboxylic acid C-1 (δ 168.6), three keto-groups C-12 (δ 207.3), C-27 (δ 201.3) and C-38 (δ 209.6) and two ester groups C-40 (δ 161.8) and C-46 (δ 174.9), with C-40 from the β-lactone ring. Furthermore, signals were observed for six olefinic methines C-2 (δ 128.5), C-3 (δ 137.8), C-4 (δ 129.1), C-5 (δ 143.2), C-28 (δ 123.0), and C-29 (δ 156.4). Finally, signals for the methyl groups C-39 (δ 29.9), C-41 (δ 8.4), C-42 (δ 9.2), C-43 (δ 14.3), C-44 (δ 19.5), C-45 (δ 17.4) and the methoxy group 8-OMe (δ 59.6) were also observed (*Supplementary information* Fig. 7). The chemical shift of C-18 (δ 101.0) indicated the presence of an acetal moiety.

COSY correlations further established five fragments including one saturated aliphatic chain. The HMBC correlations (Fig. 2) observed from H-6 and C-40 established a β-lactone moiety. The connectivity of this β-lactone to a polyene moiety was supported by the HMBC correlations between H-4 and C-6, as well as H-4 and C-40. A macrolactone bridge between H-10 and C-46 was established by HMBC correlation between them. The key HMBC correlation between H-13 and C-18 indicated the presence of an oxane ring. Moreover, H-16 and Me-43 showed HMBC correlations with C-18. Considering the number of double bond equivalents and the number of oxygen atoms inferred from HR-ESI-MS, a spiroketal structure was assigned. Finally, a planar structure with a novel carbon skeleton was proposed and named alligamycin A. The ^1^H and ^13^C NMR data, as well as the COSY and HMBC correlations of **1** are shown in *Supplementary information* Tab. 4. The complex structure exhibited no similarities to reported natural products. Meanwhile, the dispersion of the thirteen chiral centers made it challenging to elucidate the absolute configuration. Multiple attempts on crystallization were carried out on compound **1** and slow evaporation from dichloromethane/methanol solution yielded a single crystal for X-ray crystallography analysis, which successfully elucidated the configurations (*Supplementary information* Fig. 4).

**Fig. 2.**
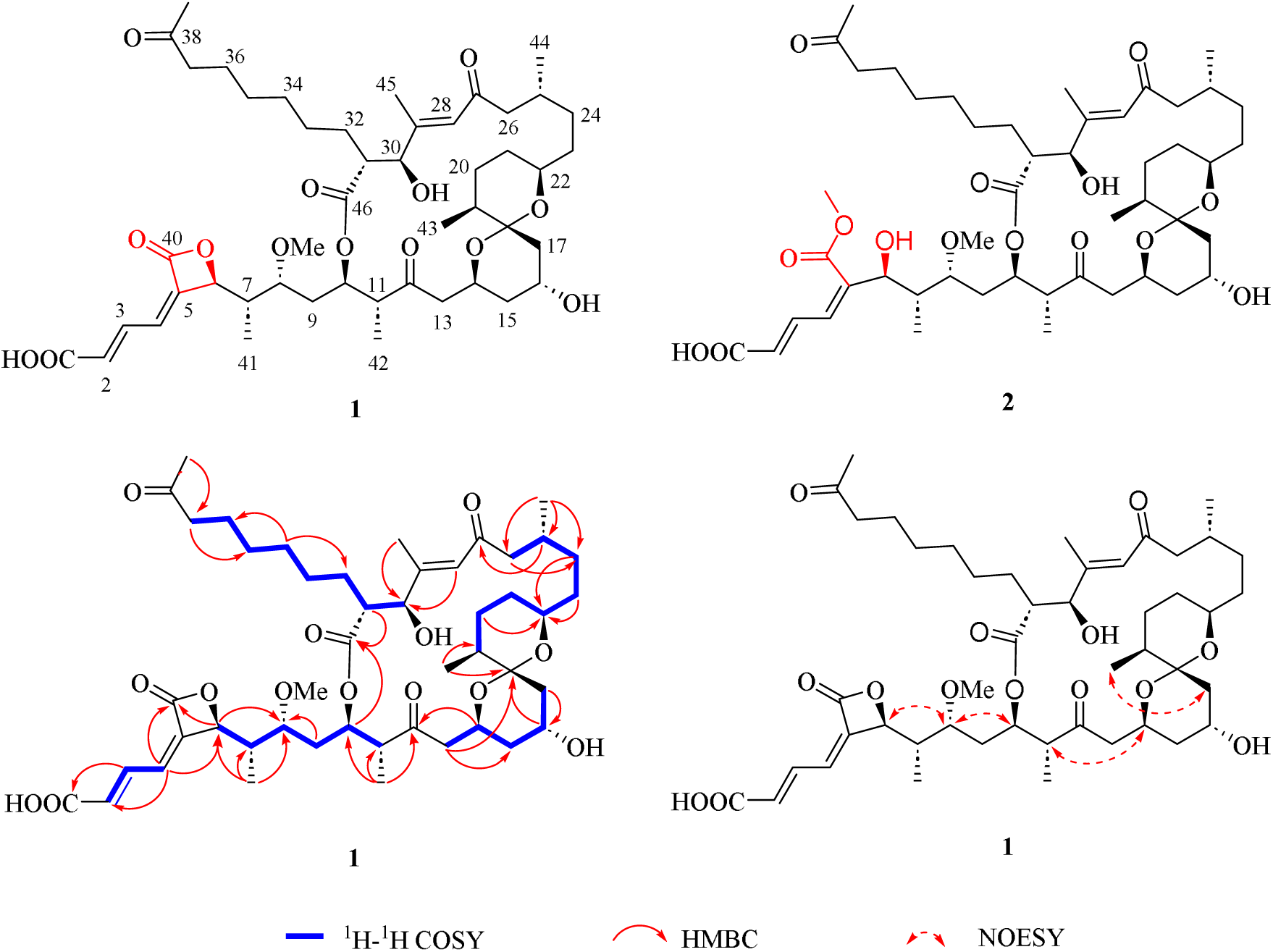
Chemical structures of alligamycin A (**1**) and B (**2**) and selected COSY, HMBC and NOESY correlations of **1**.

Alligamycin B (**2**) was assigned the molecular formula of C_48_H_74_O_15_ by HR-ESI-MS (*m/z* 889.4966 [M – H]^-^). Following 1D/2D NMR spectra, compound **2** exhibited high similarity to **1**. However, the downfield shift of the carbonyl carbon at C-40 (δ 166.9), and the presence of an additional methoxy signal (δ 3.85) showing HMBC correlation to C-40, indicating **2** consists of an opened β-lactone ring compared to **1**. Analysis of COSY, HMBC and NOESY data of **2** further confirmed the structure of alligamycin B (*Supplementary information* Fig. 14-20, Tab. 4).

### Proposed biosynthesis of alligamycins

The proposed *ali* BGC spanned 87.99 kb with 12 open reading frames (Fig. 1), including five core polyketide synthases (*aliA*, *aliC*, *aliD*, *aliE*, and *aliF*), a crotonyl-CoA carboxylase/reductase (*aliH*), an *O*-methyltransferase (*aliG*), three cytochrome P450 (*aliB*, *aliK* and *aliL*), and two hypothetical enzymes (*aliI* and *aliJ*) (*Supplementary information* Tab. 3). The core PKS genes are all transcribed in the same direction as other genes within the *ali* BGC, which consists of one loading module and fifteen extender modules. The detailed domains, ketosynthase (KS) domain, acyltransferase (AT) domain, and acyl carrier protein (ACP), with additional ketoreductase (KR), dehydratase (DH), and enoyl reductase (ER) domains and the proposed biosynthesis are shown in Fig. 3. Multiple sequence alignment revealed that most KS domains contain conserved catalytic sites, consisting of a cysteine (TA**C**SSS motif) and two histidines (EA**H**GTG and KSNIG**H**T motifs).^37^ Only the reactive cysteine in KS_1_ has been replaced by glutamine, which was observed in the platensimycin/FabF (the type II FAS homology of KS) complex.^38^ The conserved fingerprint residues for extender unit selectivity, the GHSIG and HAFH motifs are present in the nine AT domains (AT_1_, AT_2_, AT_5_, AT_7_, AT_8_, AT_9_, AT_11_, AT_12_, and AT_14_) that are specific for binding malonyl-CoA while GHSQG and YASH motifs are present in the six AT domains (AT_3_, AT_4_, AT_6_, AT_10_, AT_13_, and AT_15_) indicating binding of (*2S*)-methylmalonyl-CoA (*Supplementary information* Tab. 5 and Fig. 22).^37^ The AT_16_ from module 15, which was thought to be accountable for loading distinct extender units, exhibits an unusual IASH motif. KR domains have been previously classified as A1-, A2-, B1-, and B2-types, which reduce ketones to their L-hydroxy or D-hydroxy counterparts.^39^ We determined that the KR_3_, KR_4_, and KR_15_ appear to belong to the B1-type: all have the fingerprint LDD motif but with the absence of a P residue in the catalytic region. KR_5_ and KR_12_, with the characteristic W residue but lacking the LDD motif and H residue, were classified as A1-type KRs (*Supplementary information* Fig. 23). The other KR domains couldn’t be classified according to the sequence, but they are expected to be functional based on retrobiosynthesis of alligamycin A. Similarly, combined with structural information and analysis of conserved sites, the five DH domains (DH_6_, DH_7_, DH_8_, DH_9_ and DH_10_) were considered inactive during alligamycin biosynthesis (*Supplementary information* Fig. 24). ER_11_ is classified as L-type due to the presence of a tyrosine residue that donates a proton to the enol intermediate in L-type ER, whereas ER_13_ and ER_14_ belong to D-type due to the lack of tyrosine residue (*Supplementary information* Fig. 25). The final release and cyclization of the linear product was probably accomplished by TE domain in the last module, which contains an α/β-hydrolase catalytic core and loop regions that form a substrate-binding lid.^40^

**Fig. 3.**
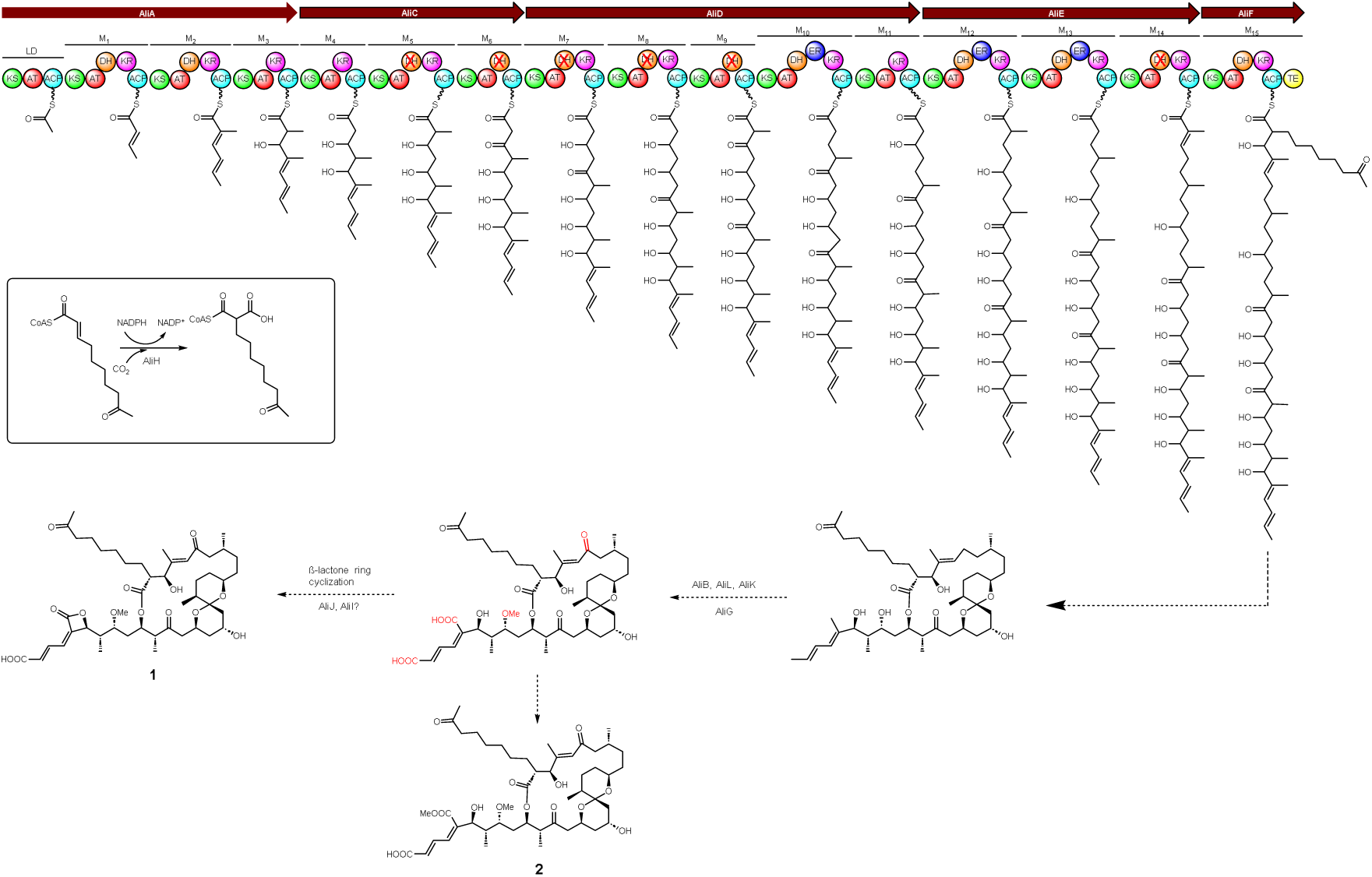
Proposed biosynthesis pathway of alligamycin A (**1**) and B (**2**). Abbreviation: KS, ketosynthase; AT, acyltransferase; DH, dehydratase; ER, enoylreductase; KR, ketoreductase; ACP, acyl carrier protein; TE, thioesterase.

In the *ali* BGC, there are three cytochromes P450 enzymes (AliB, AliK, and AliL) with conserved motifs of this family (*Supplementary information* Fig. 26).^41^ The cytochrome P_450_ enzyme BonL from *Burkholderia gladioli* was previously identified to confer the C-22 carboxyl group in the biosynthesis of bongkrekic acid via sequential six-electron-oxidation.^42^ In addition, there are some cytochromes P_450_ enzymes that catalyze the carboxyl group formation in microbial SMs, such as XiaM, PimG, AmphN, NysN, and FscP.^43–45^ To identify the cytochromes P450s responsible for carboxylation, we built the Hidden Markov model (HMM) based on these sequences and the results showed that AliL exhibited the closest match to these known P450s with an E-value of 4e-117. In addition, bioinformatic analysis of XiaM indicated that the P450s with carboxylation function harbor highly conserved segment AGHET, which was also presented in AliK (*Supplementary information* Fig. 26). Therefore, AliK and AliL, were considered putative candidates for catalyzing the two-step oxidation to form carboxyl groups at positions C-1 and C-5 during alligamycin A biosynthesis. AliB, homologous cytochrome P450 monooxygenase CftA in the clifednamide biosynthetic gene cluster with 45.6% similarity,^46^ was likely to accomplish the oxidation at C-27 in alligamycin A. We constructed mutant strains of *S. iranensis* inactivating the AliB, AliK and AliL enzymes (designated as Δ*aliB*, Δ*aliK*, and Δ*aliL*) (*Supplementary information* Fig. 27). Despite our efforts, no expected intermediates were observed in either mutant, likely due to their low yield.

Currently, there are only few enzymatic mechanisms corresponding to β-lactone ring formation that have been reported.^47^ For example, (1) the intramolecular cyclization from seven-membered ring, catalyzed by cyclase VibC in vibralactone biosynthesis;^48^ (2) the tandem aldol-lactonization bicyclization reaction to generate the γ-lactam-β-lactone structure, catalyzed by standalone ketosynthase SalC in salinosporamide A biosynthesis;^49^ (3) the β-lactone formation during the intramolecular attack of the *β*-hydroxyl group onto the thioester carbonyl, catalyzed by the C-terminal TE domain of ObiF in obafluorin biosynthesis or esterase GloD in globilactone A biosynthesis;^50,51^ (4) the conversion of β-hydroxyl to β-lactone, catalyzed by β-lactone synthase OleC in olefin biosynthesis.^52,53^ We proposed that the β-lactone formation in alligamycin A involves a two-step oxidation to form a carboxylic acid, followed by dehydration. It is likely that either the cytochrome P450 enzyme AliK or AliL is involved in the process, and we also identified several genes with unknown functions, such as *aliJ* and *aliI*. AliJ was annotated as a member of UbiD family decarboxylases, but it exhibits very distinguished divergences from other proteins in this family based on sequence similarity network (*Supplementary information* Fig. 28). In addition, NCBI Blastp results showed that proteins homologous to AliI have not been reported to have any clear biological functions. Inactivation of both genes abolished the production of alligamycin A (Supplementary information Fig. 27).

Phylogenetic analysis indicated that AliH is a separate branch and clusters with other crotonyl-CoA reductase/carboxylases (CCRs) that catalyze long-chain precursors like butylmalonyl-CoA and hexylmalonyl-CoA.^54,55^ The previous study showed that the large residue Phe380 in the CCR from *S. coelicolor* may constrain the potential pocket to accept long-chain substrates, which were also found in most ethylmalonyl-CoA specific crotonyl-CoA reductase/carboxylases, while its corresponding residue in AliH is smaller cysteine (*Supplementary information* Fig. 29).^56^ Hence, we proposed that AliH catalyzes the conversion leading to the formation of 7-oxo-octylmalonyl-CoA in alligamycin A, which is likely synthesized from a precursor that has previously been isolated from other *Streptomyces* species.^57^ AliG, an O-methyltransferase homologous to AveD in avermectin biosynthesis (40% identity and 55% similarity),^58^ was proposed to catalyze the methylation of a hydroxyl group in C-8 of alligamycin A.

### Antifungal and cytotoxic assays of alligamycins

Microbial β-lactone natural products are chemically diverse and have been employed in antimicrobial, antiviral, anticancer, and antiobesity therapeutics.^47^ In the initial antifungal assay against *A. niger* ATCC 1015, alligamycin A exhibited the strongest antifungal effect, while alligamycin B with an opened β-lactone ring did not show antifungal activity (MIC >50 µg/mL). Since both alligamycins contain an α,β-unsaturated ester, this suggests that the electrophilic nature of the β-lactone (and not the ability to act as a Michael acceptor) was responsible for the antifungal activity of alligamycin A. The *in vitro* antifungal susceptibility screens of alligamycin A was further performed against 34 fungal species, represented by 38 clinical isolates, in comparison to antifungal drugs using a microdilution assay according to the EUCAST protocol. Satisfyingly, alligamycin A demonstrated potent antifungal effects against several sections of the genus *Aspergillus* and the genus *Talaromyces* with minimal inhibitory concentration (MIC) ranging from 0.06 to 8 µg/mL (Tab. 1, *Supplementary information* Tab. 6). The most potent activity of alligamycin A compared to standard drugs were found in in the *Aspergillus* section *Terrei* with high MICs for amphotericin B and in the section *Usti* with intrinsically high MICs for azoles. In *Aspergillus* sections *Flavi*, *Nidulantes*, and *Nigri*a, as well as in the section *Talaromyces*, a distinctly changed morphology of the hyphae was found for all species but low MICs were observed only in some species. Only a weak effect was observed for the two *Fusarium* species studied. *Aspergillus* species belong to the section *Fumigati*, *Candida* spp., *Scedosporium prolificans*, *Purpureocillium lilacinum*, *Trichoderma longibrachiatum* and *Mucorales* were unaffected by alligamycin A (*Supplementary information* Tab. 6). Importantly, alligamycin A did not exhibit cytotoxicity against the human acute promyelocytic leukemia cell line (HL-60) with IC_50_ >10 µg/mL highlighting an important advantage over other anti-mycotics and its potential for drug development.

**Tab. 1.** Antifungal activities of alligamycin A (**1**) in comparison with amphotericin B, itraconazole and voriconazole, the values for MIC (MEC) are in mg/L. MIC: Minimum Inhibition Concentration; MEC: Minimum Effective Concentration.

| Species | Section | MIC (MEC) in mg/L |  |  |  |
| --- | --- | --- | --- | --- | --- |
|  |  | amphotericin B | itraconazole | voriconazole M | alligamycin A |
| *A. brasiliensis* | *Nigri* | 0.5 | 2 | 2 | 0.5 (0.25) |
| *A. luchuensis* | *Nigri* | 0.25 | n.a. | 1 | 0.5 (0.25) |
| *A. citrinoterreus* | *Terrei* | 4 | 0.5 | 0.5 | 0.5 (0.25) |
| *A. neoafrikanus* | *Terrei* | 2 | 0.5 | 0.5 | 0.5 (0.25) |
| *A. terreus* | *Terrei* | 8 | 0.5 | 1 | 0.125 (0.125) |
| *A. calidoustus* | *Usti* | 16 | 8 | 4 | 0.5 (0.25) |
| *A. ustus* | *Usti* | 1 | >8 | >8 | 0.125 (0.06) |
| *T. marneffeii* | | 0.06 | $\leq 0.016$ | $\leq 0.016$ | 0.125 (0.125) |
| *T. purpureogenes* |  | 1 | >8 | 8 | 0.5 (0.25) |

### Studies of the mechanism of action of alligamycin A

To obtain initial insights into the alligamycin mode of action, we carried out a label-free quantitative proteomics analysis of the model organism *A. niger* ATCC 1015. We initially cultured *A. niger* in PDB for 16 hours, followed by 1-hour and 4-hour treatments with alligamycin A. The proteomics analysis showed a total of 4,447 proteins (40.4% of the total encoded proteins) and 3,992 of these proteins (89.8% of the total detected proteins) did not show significant differences between the treatments (1-hour or 4-hour treatment with alligamycin A) and the control (without alligamycin A treatment). Compared with the control, the abundance of 134 proteins significantly increased, while 60 proteins significantly decreased after 1-hour alligamycin A treatment. In comparison, the abundance of 273 proteins significantly increased and 113 proteins significantly decreased after 4-hour alligamycin A treatment (Fig. 4). The Cluster of Orthologous Groups (COGs) functional classification of these differential proteins was performed by eggNOG-mapper v2.^59^ Except for (S) function unknown, these proteins were mainly clustered into a few biological processes including (Q) Secondary structure, (C) Energy production and conversion, (E) Amino acid metabolism and transport, and (O) Post-translational modifications, protein turnover, chaperone functions (*Supplementary information* Fig. 30). The Gene Ontology (GO) annotation of these proteins was mainly involved in the metabolic process, cellular process, and single-organism process during the biological process (*Supplementary information* Fig. 31). Cellular component analysis showed that these proteins were mainly localized intracellularly, especially in organelles. Meanwhile, most of these proteins participated in catalytic activity and binding in the category of molecular function. The Kyoto Encyclopedia of Genes and Genomes (KEGG) pathway enrichment analysis showed that these proteins mainly participate in amino acids metabolism and purine metabolism (*Supplementary information* Fig. 32).

**Fig. 4.**
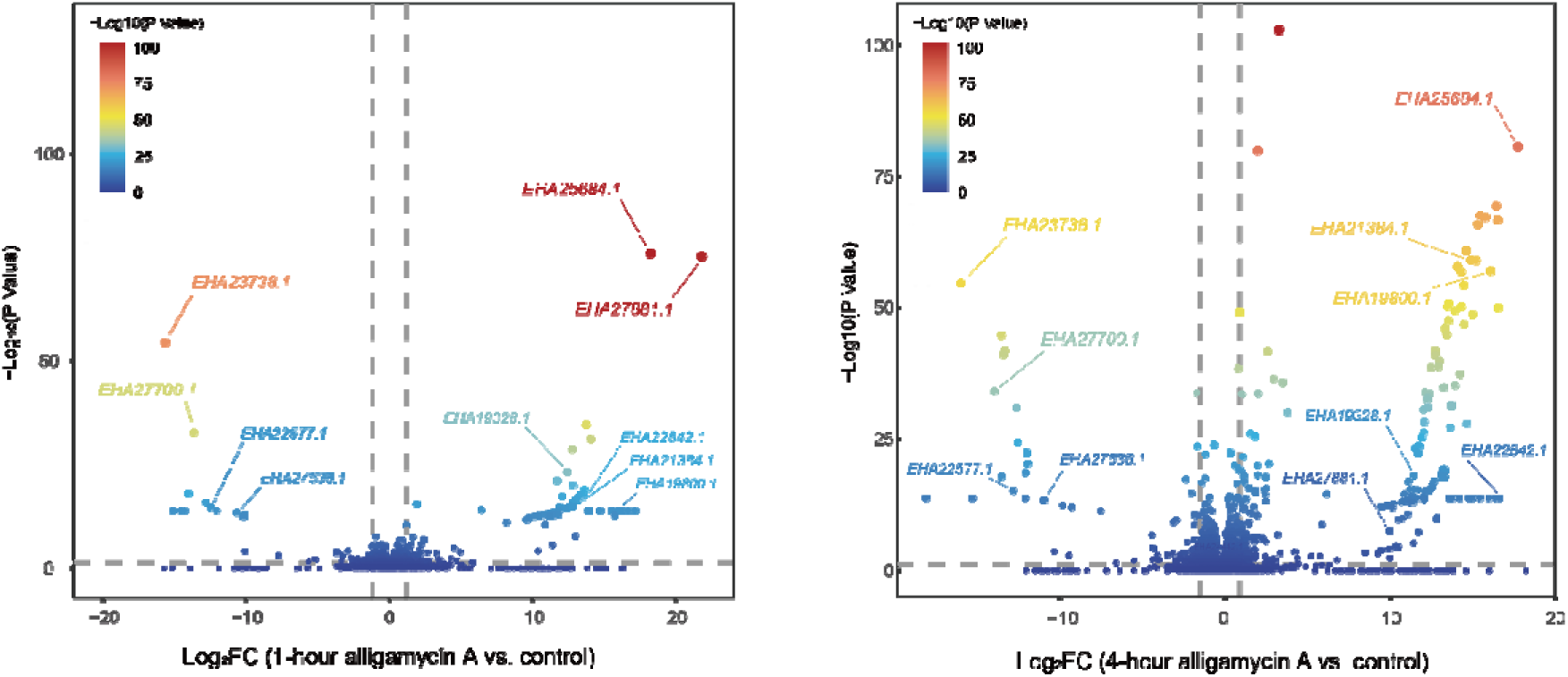
Volcano plot of the protein abundance of *A. niger* after 1-hour and 4-hour alligamycin A treatments *versus* control based on proteomics analysis.

Proteomic changes in fungi induced by known antifungal agents with well-described targets were subsequently studied. The abundance of Erg13 (EHA22977.1, involved in ergosterol biosynthesis) and manganese-superoxide dismutase (EHA27012.1, involved in oxidative stress response), which was regarded as polyene antibiotic (such as amphotericin B) responsive protein,^60^ was not significantly altered in the presence of alligamycin A. Allylamine antibiotics (naftifine and terbinafine) inhibit squalene epoxidase, causing the accumulation of squalene, and thereby damaging the intracellular membranes of fungi.^61^ Alligamycin A does not belong to the allylamine family and we did not observe any changes in the squalene epoxidase (EHA21184.1). Azoles are synthetic antifungal agents derived from non-natural origin that inhibit cytochrome P450 sterol-14*α*-demethylase (CYP51) to block the ergosterol biosynthesis in the fungi.^62^ Similarly, alligamycin A lacked the azoles substructure and we did not observe impact on the abundance of CYP51 (EHA19435.1). Natural echinocandins comprise cyclic hexapeptide and lipid residues, like caspofungin, anidulafungin, and micafungin, preventing biosynthesis of glucans of the fungal cell wall through non-competitive inhibition of 1,3-β-D-glucan synthase.^63^ Previous proteomics analyses had proven that the level of chitinase ChiA1 in *A. fumigatus* was significantly decreased in response to caspofungin.^64^ It may represent a common effect of self-resistance development due to caspofungin increased chitin content via induction of chitin synthases during cell wall remodeling.^65^ We did not observe the abundance of 1,3-β-D-glucan synthase (EHA18547.1) and ChiA1-like chitinase (EHA28582.1) altered significantly after 1-hour and 4-hour treatment of alligamycin A in comparison to the control group.

The fungal cell wall constitutes a critical structural component that maintains cell shape, protects fungi against environmental stress, and plays roles in growth, invading ecological niches invasion, and counteracting the host immune response.^66^ Fungal cell walls are composed mainly of glucans, chitin, and glycoproteins, synthesized by glycosyltransferases, glycoside hydrolases, and transglycosylases.^67^ Congo Red Hypersensitivity (CRH) family transglycosylases are usually highly expressed in multiple stages during the conidial germination of fungi and may be involved in cell wall synthesis and stability. The volcano plot depicting differential proteins indicated that a CRH family transglycosylase (EHA23738.1) was significantly inhibited, suggesting alligamycin A might interfere with the formation of the fungal cell wall. Trehalose-6-phosphate synthase/phosphatase is another important enzyme for cell wall integrity and fungal virulence in various *Aspergillus* species.^68^ The relative abundance of trehalose 6-phosphate synthase (EHA27700.1) was also significantly decreased after alligamycin A treatment. The above data indicated that the mode of action of alligamycin A could be through disruption of fungal cell wall biosynthesis and stability (Fig. 4). Besides, we found that the two enzymes Erg24 (EHA26587.1) and Erg27 (EHA20711.1), which catalyze the biosynthesis of 4,4-dimethylcholesta-8,14,24-trienol to fecosterol,^69^ were remarkably upregulated after alligamycin A treatment (Fig. 4). Drug efflux and resistance in fungi could be mediated by ATP-binding cassette (ABC) transporters, such as CDR1 was induced to high expression in *A. fumigatus* AF293 in the presence of azole antibiotics.^70^ The expression of an ABC transporter (EHA25684.1) was activated after alligamycin A treatment. We noted significant alterations in the abundance of several putative alligamycin A-responsive proteins. For example, the relative abundance of several proteins including inositol hexaphosphate kinase KCS1 (EHA27881.1), ubiquitin-protein ligase ASI3 (EHA19328.1), mitochondrial phosphate carrier protein PIC2 (EHA22842.1), pyridoxamine 5’-phosphate oxidase (EHA19800.1) and glucohydrolase (EHA21384.1) was significantly increased, but oxygen-dependent FAD-linked oxidoreductase (EHA22577.1) and dienelactone hydrolase (EHA27638.1) was significantly decreased in alligamycin A-treated samples. Intriguingly, a number of proteins of unknown function displayed variations in abundance attributable to alligamycin A and their specific roles warrant further investigation.

## Conclusion

In this study, we report the discovery of a novel class of antifungal agent alligamycin A, featuring an unprecedented carbon skeleton. Alligamycin A exhibits selective antifungal activities against several pathogenic *Aspergillus* species, some of which are intrinsically resistant to amphotericin B or azoles, as well as against *Talaromyces* species. We confirmed the BGC of alligamycin A through a diverse array of *in silico* and *in vivo* analyses. Proteomics data indicated that treatment with alligamycin A results in the inhibition of transglycosylase and trehalose-6-phosphate synthase/phosphatase, thereby suggesting its potential role in targeting fungal cell wall biosynthesis.

## Methods and Materials

### Strains, plasmids, and culture conditions

All strains and plasmids used in this study were summarized in *Supplementary information* Tab. 7. All primers used in this study were summarized in *Supplementary information* Tab. 8. All constructed *E. coli* strains were grown in lysogeny broth (LB) liquid or on agar medium at 37 °C. Wild-type *Streptomyces* strains and its mutants were cultivated on Mannitol Soya Flour (MS) agar medium (20.0 g mannitol, 20.0 g soya flour, 20.0 g agar, 1.0 L tap water, pH=7.0-7.5) at 30 □. The small-scale fermentation of *Streptomyces* strains was performed in 250 mL flask using 50 mL MS liquid medium, shaking at 200 rpm, 30 □ for 7 days. The large-scale fermentation of *Streptomyces* strains were carried out in medium 2 (CaCl_2_·2H_2_O, 3.0 g; citric acid/Fe III, 1.0 g; MnSO_4_·H_2_O, 0.2 g; ZnCl_2_, 0.1 g; CuSO_4_·5H_2_O, 0.025 g; Na_2_B_4_O_7_·10H_2_O, 0.02 g; Na_2_MoO_4_·2H_2_O, 0.01 g; and oatmeal, 20.0 g, in 1.0 L distilled water), at 200 L scale in a 300 L fermentation vessel, for 6 days with aeration of 25-50 L min^-1^, stirring at 200 rpm, 28 °C, with a pH range of 5.4-6.4. Antibiotics such as apramycin (50 mg mL^-1^), kanamycin (50 mg mL^-1^) or chloramphenicol (25 mg mL^-1^) were appropriately used for resistance selection.

### Genomic DNA extraction, sequencing, and assembly

The *S. iranensis* culture was grown in 50 mL sterilized liquid ISP2 medium (yeast extract 4.0 g; malt extract 10.0 g; and dextrose 4.0 g in 1.0 L distilled water, pH = 7.2) in 250 mL flask at 30 °C and 160 rpm for 5 days to generate sufficient biomass. The genomic DNA of *S. iranensis* was isolated using the QIAGEN Genomic-tip 20/G kit with a modified protocol and the cell pellet was ground using a mortar and pestle submerged in liquid nitrogen.^71^ The genomic DNA sequencing was performed combining Oxford Nanopore Technologies MinION and Illumina MiSeq systems. To generate the nanopore data, both the SQK-RBK004 and the SKQ-RBK110-96 rapid barcoding kits were used for construction of three libraries, which were sequenced on two separate 9.4.1 flow cells. The *de novo* assembly was performed using Flye (v2.9-b1768) with the parameters --nano-hq --t 12. Since the genome could not be fully resolved using existing assemblers, the terminal inverted repeat edge from the unambiguous assembly repeat graph were manually attached to both ends of the chromosome edge and the orientations were verified by mapping of the reads on the new assembly and manual inspection. This approach was later suggested by the Flye assembler Misha. The Unicycler (v0.4.8) polishing module was used to polish the nanopore assembly with Illumina data.

### Genetic manipulation

To verify the BGC of alligamycin and its individual biosynthetic genes, the classical CRISPR-Cas9 method, and advanced CRISPR-cBEST base editing toolkit were used to construct gene-inactive mutants.^36^ The function-specific plasmids (*Supplementary information* Tab. 7) were constructed according to the respective protocol followed by introducing into wild-type *S. iranensis* HM 35 by conjugation with donor strain ET12567/pUZ8002 on SFM solid medium (soya flour 20.0 g; mannitol 20.0 g; and bacteria agar 20.0 g, in 1.0 L distilled water, pH = 7.2) according to modified protocol. The exconjugants after resistance screening (50 μg mL^-1^ apramycin and 25 μg mL^-1^ nalidixic acid) were further verified by DNA extraction, PCR reaction, and Sanger sequencing.

DNA polymerases (Q5^®^ High-Fidelity 2X Master Mix with Standard Buffer and OneTaq^®^ 2X Master Mix with Standard Buffer) and restriction enzymes (NcoI, EcoRI) were purchased from New England Biolabs. PCR amplifications and restriction enzyme digestions were carried out on Bio-Rad’s thermal cyclers according to the manufacturer’s instructions. Plasmid DNA extraction was performed using NucleoSpin Plasmid EasyPure Kit (Macherey-Nagel, Germany). DNA purification was conducted on 1% Tris-acetate-EDTA (TAE) agarose gel followed by using NucleoSpin Gel and PCR Clean-up Kits (Macherey-Nagel, Germany). One Shot™ Mach1™ T1 Phage-Resistant Chemically Competent *E. coli* from Invitrogen™ was used for transformation. All oligonucleotides were ordered from Integrated DNA Technologies (*Supplementary information* Tab. 8) and Sanger sequencing was offered by Eurofins Genomics (Luxembourg).

### HR-ESI-MS analysis

HR-ESI-MS data were acquired on an Agilent Infinity 1290 UHPLC system equipped with a diode array detector and coupled to an Agilent 6545 QTOF MS equipped with Agilent Dual Jet Stream ESI. Separation was achieved on a 250 × 2.1 mm i.d., 2.7 μm, Poroshell 120 Phenyl Hexyl column (Agilent Technologies) held at 40 °C. The sample, 1 μL, was eluted at a flow rate of 0.35 mL min^−1^ using a linear gradient from 10% acetonitrile/water buffered with 20 mM formic acid to 100% acetonitrile in 15 min, held for 2 min and equilibrated back to 10% acetonitrile/water in 0.1 min. Starting conditions were held for 3 min before the following run. The MS settings were as follows: drying gas temperature of 160 °C, a gas flow of 13 L min^−1^, the sheath gas temperature of 300 °C and flow of 16 L min^−1^. Capillary voltage was set to 4000 V and nozzle voltage to 500 V in positive mode. All data were processed using Agilent MassHunter Qualitative Analysis software (Agilent Technologies, USA). All solvents used for chromatography and HR-MS were purchased from VWR Chemicals with LC-MS grade, while for metabolites extraction, the solvents were of HPLC grade.

### Fermentation and isolation

The culture broth from large-scale fermentation (described above) was filtered and loaded onto an Amberchrom 161c resin LC column (200 × 20 cm, 6 L). Elution with a linear gradient of H_2_O-MeOH (from 30% to 100% v/v, flow rate 0.5 L min^-1^, in 58 min) afforded seven fractions (Fr.A-Fr.G). Fr.G was first fractionated by silica gel chromatography with a CH_2_Cl_2_-CH_3_OH solvent system to yield 16 fractions, Fr.1-Fr.16. Fr.7 was further separated by a Sephadex LH-20 (MeOH) column, and twelve sub-fractions were obtained. The sub-fractions were separated by semipreparative HPLC RP-C_18_ using MeCN-H_2_O as solvent system to afford compound **1** and **2.**

Alligamycin A (**1**): white solid;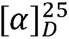15 (0.1 mg/mL, CH_3_OH), UV/vis (CH_3_CN/H_2_O) λ_max_ 230, 270 nm; ECD λ_ext_ (Δε) (CH_3_OH) 235 (–9.78), 267 (7.02), 301 (–3.31) nm; IR (ATR) *v*_max_ 2932, 2748, 2704, 1810, 1713, 1686, 1619, 1457, 1382, 1173, 1134, 1085, 1057, 1013, 991, 947 cm^-1^; (+)-HR-ESI-MS *m/z* 881.4659 [M + Na]^+^ (calcd for C_47_H_70_O_14_, 881.4658).

Alligamycin B (**2**): white solid; 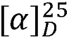 18 (0.2 mg/mL, CH_3_OH); UV/vis (CH_3_CN/H_2_O) λ_max_ 230, 270 nm; IR (ATR) *v*_max_ 2990, 2942, 2919, 2831, 2035, 1449, 1416, 1119, 1022 cm^-1^; (-)-HR-ESI-MS *m/z* 889.4971 [M - H]^-^ (calcd for C_48_H_74_O_15_, 889.4955).

### NMR spectroscopy

NMR spectra were recorded on 800 MHz Bruker Avance III spectrometer equipped with a TCI CryoProbe using standard pulse sequences. Measurements were carried out using ^1^H and ^13^C NMR, ^1^H-^13^C heteronuclear single quantum coherence (HSQC), ^1^H-^13^C heteronuclear multiple bond correlation (HMBC), ^1^H-^1^H correlation spectroscopy (COSY), ^1^H-^1^H nuclear overhauser effect spectroscopy (NOESY). Chemical shifts (δ) were reported in parts per million (ppm) and the ^1^H and ^13^C NMR chemical shifts were referenced to the residual solvent peaks at δ_H_ 7.26 and δ_C_ 77.16 ppm for CDCl_3_. Data are described as follows: chemical shift, multiplicity (*br* = broad, *s* = singlet, *d* = doublet, *t* = triplet, *dd* = doublet of doublet, *m* = multiplet and *ov* = overlapped) and coupling constants (in Hertz). All NMR data were processed using MestReNova 14.0.

### Crystal Data for alligamycin A

The suitable crystal was selected and mounted in a nylon loop directly from the ethanol suspension and frozen in liquid nitrogen on a Synchrotron diffractometer. X-ray data collection of **1** was performed on an Agilent Supernova Diffractometer using CuKα radiation, and the crystal was kept at 100 K during data collection. Using Olex2,^72^ the structure was solved with the XT^73^ structure solution program using Intrinsic Phasing and refined with the SHELXL^74^ refinement package using Least Squares minimization. Crystal data for C_47_H_70_O_14_ (*M* =859.03 g/mol): monoclinic, space group P2_1_ (no. 4), *a* = 10.118(7) Å, *b* = 16.799(7) Å, *c* = 14.081(5) Å, β = 103.385(12)°, *V* = 2328(2) Å^3^, *Z* = 2, *T* = 100 K, μ (synchrotron) = 0.089 mm^-1^, *Dcalc* = 1.225 g/cm^3^, 17409 reflections measured (3.836° ≤ 2*θ* ≤ 45.2°), 5762 unique (*R*_int_ = 0.0646, *R*_sigma_ = 0.0641) which were used in all calculations. The final *R*_1_ was 0.0646 (I > 2σ(I)) and *wR*_2_ was 0.1523 (all data).

### Sample preparation for proteomics analysis

Approximately 1.0 × 10^7^ conidia mL ^-1^ of *A. niger* ATCC 1015 strain were used to inoculate in 500 mL flasks containing 100 mL of liquid cultures (PDB) (three biological replicates) and were incubated in a reciprocal shaker at 28 □ in 180 rpm for 16 hours. Afterwards, samples in the treatment group were treated with 0.2 μg (concentration of 2 ng/mL) of alligamycin A for 1 hour and 4 hours. The mycelia were then harvested by filtering, washed thoroughly with sterile water and quickly frozen in liquid nitrogen. Pellets were lysed in 100 µL lysis buffer (6 M guanidium hydrochloride, 10 mM TCEP, 40 mM CAA, 50 mM HEPES, pH 8.5) by boiling the samples at 95 □ for 5 mins, followed by sonicating on high for 5 × 60 sec on/30sec off using the Bioruptor Pico sonication water bath (Diagenode) and lastly disrupting twice with the TissueLyser (QIAGEN) going from 3 to 30 Hz in 1 min. Lysed samples were centrifuged at 18,000 g for 10 mins, and supernatants were transferred to clean LoBind Eppendorf tubes. Protein concentration was determined by BCA rapid gold (Thermo) and 10 µg of protein was taken forward for digestion. Samples were diluted 1:3 with digestion buffer (10% acetonitrile in 50mM HEPES pH 8.5) and incubated with 1:100 enzyme to protein ratio of LysC (MS Grade, Wako) at 37 □ for 4 hours. Samples were further diluted to a final 1:10 with more digestion buffer and digested with 1:100 trypsin for 18 hours at 37 □. After digestion, samples were acidified with TFA and desalted using the SOLAµ^TM^ SPE plate (HRP, Thermo).^75^ Between each application, the solvents were spun through by centrifugation at 1,500 rpm. For each sample, the filters were activated with 200 µL of 100% Methanol, then 200 µL of 80% Acetonitrile, 0.1% formic acid. The filters were subsequently equilibrated 2× with 200 µL of 1% TFA, 3% acetonitrile, after which the sample was loaded. After washing the tips twice with 200 µL of 0.1% formic acid, the peptides were eluted into clean 0.5 mL Eppendorf tubes using 40% acetonitrile, 0.1% formic acid. The eluted peptides were concentrated in an Eppendorf Speedvac. Samples were reconstituted in 12 µL A* buffer with iRT peptides (Biognosys).

### MS analysis for proteomics analysis

Peptides were loaded onto a 2 cm C_18_ trap column (ThermoFisher 164946), connected in-line to a 15 cm C_18_ reverse-phase analytical column (Thermo EasySpray ES904) using 100% Buffer A (0.1% Formic acid in water) at 750 bar, using the Thermo EasyLC 1200 HPLC system, and the column oven operating at 30°C. Peptides were eluted over a 70 minutes gradient ranging from 10% to 60% of Buffer B (80% acetonitrile, 0.1% formic acid) at 250 µL/min, and the Orbitrap Exploris instrument (ThermoFisher Scientific) was run in DIA mode with FAIMS ProTM Interface (ThermoFisher Scientific) with CV of -45 V. Full MS spectra were collected at a resolution of 120,000, with an AGC target of 300% or maximum injection time set to ‘auto’ and a scan range of 400–1000 m/z. The MS2 spectra were obtained in DIA mode in the orbitrap operating at a resolution of 60.000, with an AGC target 1000% or maximum injection time set to ‘auto’, a normalised HCD collision energy of 32. The isolation window was set to 6 m/z with a 1 m/z overlap and window placement on. Each DIA experiment covered a range of 200 m/z resulting in three DIA experiments (400-600 m/z, 600-800 m/z and 800-1000 *m/z*). Between the DIA experiments a full MS scan is performed. MS performance was verified for consistency by running complex cell lysate quality control standards, and chromatography was monitored to check for reproducibility.

### Data process for proteomics analysis

The raw files were analyzed using Spectronaut^TM^ (version 17.4) spectra were matched against the reviewed *A. niger* ATCC 1015 NCBI database. Dynamic modifications were set as Oxidation (M) and Acetyl on protein N-termini. Cysteine carbamidomethyl was set as a static modification. All results were filtered to a 1% FDR, and protein quantitation done on the MS1 level. Only proteotypic peptides were used for quantification and protein groups were inferred by IDPicker.

### Antifungal susceptibility testing

In vitro antifungal susceptibility of alligamycin A against 34 fungal species, represented by 38 clinical isolates, was tested in comparison to approved antifungal drugs amphotericin B (AMB; European Pharmacopoeia, Strasbourg, France); itraconazole (ITZ), and voriconazole (VCZ; Pfizer Inc., Peapack, NJ, USA) using broth microdilution technique following the European Committee on Antimicrobial Susceptibility Testing (EUCAST) standard methodology for yeasts or filamentous fungi respectively. In contrast to the EUCAST protocol, microdilution plates were prepared by 2-fold serial dilutions of the antifungal agents. Filamentous fungi were grown on malt extract agar (MEA) for 2 to 7 days at 35 °C and yeasts were cultivated on yeast extract peptone dextrose agar (YPD) for 24 hours. Spore or yeast cell suspensions were counted with a hemocytometer. Minimum inhibitory concentrations (MIC) endpoints of filamentous fungi were defined as 100% reduction in growth and were determined visually using a mirror after 48 h of incubation at 35 °C. Microdilution plates of yeasts were read with a microdilution plate reader (Infinite® M Nano plus, Tecan) and MIC endpoints were defined as the lowest drug concentration giving inhibition of growth of ≥ 50% of that of the drug-free control. Since the mode of action of alligamycin A is unknown, the minimum effective concentration (MEC) was determined as well by reading the microplates with the aid of an inverted microscope (Eclipse Ts2, Nikon). *A. fumigatus* ATCC 204305 and *Candida parapsilosis* ATCC 22019 were used as reference strains.^76,77^

### Cytotoxic activities of alligamycins

Cytotoxicity against the human cell line HL-60 was evaluated using Alamar Blue (Thermos Scientific, Kansas, USA). The assay (Hamid et al. 2004) was performed in 96 well plates (Costar 3595, Corning, New York, USA), with an assay volume of 200 µL. The software Prism 5.03 was used for data analysis (GraphPad Software, USA).^78^

## Data availability

The genome sequence of *S. iranensis* HM35 has been deposited in the NCBI under BioProject accession number PRJNA1026072 and BioSample accession number SAMN37733152. The genome sequence could be access under Genbank accession number CP136563. The crystal data of alligamycin A was deposited in CCDC database (https://www.ccdc.cam.ac.uk/) with accession number 2178562. The metabolomics data of wild-type and mutant strains has been deposited in the MassIVE repository (https://massive.ucsd.edu/) under identifier MSV 000094394. The mass spectrometry for proteomics analysis have been deposited in the ProteomeXchange Consortium (https://www.proteomexchange.org/) with data set identifier MassIVE MSV000094235. The NMR data were deposited at NP-MRD, with the accession number of NP0332733 and NP0333731, for alligamycin A and B, respectively.

## Acknowledgements

L.D. acknowledges funding from DTU Skylab, Novo Nordisk Foundation (NNF23OC0082881) and Innovation Fund Denmark (Grant 3141-00013A). M.H.C. acknowledges funding for the DTU Screening Core from the Novo Nordisk Foundation (NNF19OC0055818) and the Carlsberg Foundation (CF19-0072). Y.Q. acknowledges funding from Novo Nordisk Foundation (NNF22OC0079928). T.W. acknowledges funding from the Novo Nordisk Foundation (NNF20CC0035580). The NMR Center DTU and the Villum Foundation are acknowledged for access to the 800 MHz spectrometer. The metabolomic data was generated at DTU Metabolomics Core facilities with the help of Dr. A. Andersen. Thanks to Prof. M. Shaaban for repurification of alligamycin A and recording the ECD spectrum. Thanks to S. Schmidt for helping with isolation of alligamycin B. Thanks to Prof. Jacob Blæsbjerg Hoof for the *A. niger* strain and advice.

## Competing interests

The authors declare that there are no competing interests. Two patent applications were filed with the numbers of PCT/EP2023/067953 and EP24150176.6.

## Additional Information

**Supplementary information:** The online version contains supplementary material available at xxx.

**Correspondence and requests** for materials should be addressed to Ling Ding.

## Notes

### Summary of Updates

The compound name is revised to alligamycin. We added one new analogue to demonstrate the mode of action.

