## Supplemental data for "Alligamycin A, an unprecedented antifungal β-lactone spiroketal macrolide from *Streptomyces iranensis*"

[#] Contribute equally

### Table of Contents


| Table and Figures | Page |
|---|---|
| **Tab. 1.** All detected BGCs belonging to the GCF_00315 in BiG-FAM database. | 4 |
| **Tab. 2.** The biosynthetic gene clusters in *S. iranensis* based on genome resequencing. | 5 |
| **Tab. 3.** The function annotation of biosynthetic genes in *ali* BGC. | 7 |
| **Tab. 4.** $^{1}$H (800 MHz) and $^{13}$C (200 MHz) NMR data of **1** and **2** ($CDCl_3$). | 8 |
| **Tab. 5.** Alignment of conserved motifs in the active site of AT domains. | 11 |
| **Tab. 6.** Antifungal assays of alligamycin A. | 12 |
| **Tab. 7.** The strains and plasmids used in this study. | 14 |
| **Tab. 8.** The primers used in this study. | 15 |
| **Fig. 1.** Co-culture of *S. iranensis* and *Aspergillus* on PDB agar medium. | 17 |
| **Fig. 2.** Domain annotation of *ali* BGC in *S. iranensis* before and after whole-genome resequencing. | 18 |
| **Fig. 3.** The biosynthetic gene architecture of alligamycin A and other BGCs from GCF_00315. | 19 |
| **Fig. 4.** ORTEP drawing of compound **1**. | 20 |
| **Fig. 5.** HR-ESI-MS spectrum and UV spectrum of **1**. | 20 |
| **Fig. 6.** $^{1}$H NMR (800 MHz) spectrum of **1** in $CDCl_3$**.** | 21 |
| **Fig. 7.** $^{13}$C NMR (200 MHz) spectrum of **1** in $CDCl_3$. | 21 |
| **Fig. 8.** HSQC (800 MHz) spectrum of **1** in $CDCl_3$. | 22 |
| **Fig. 9.** $^{1}$H-$^{1}$H COSY (800 MHz) spectrum of **1** in $CDCl_{3.}$ | 22 |
| **Fig. 10.** HMBC (800 MHz) spectrum of **1** in $CDCl_3$. | 23 |
| **Fig. 11.** NOESY (800 MHz) spectrum of **1** in $CDCl_3$. | 23 |
| **Fig. 12.** ECD spectrum of **1**. | 24 |
| **Fig. 13.** IR spectrum of **1**. | 24 |
| **Fig. 14.** HR-ESI-MS spectrum and UV spectrum of **2**. | 25 |
| **Fig. 15.** $^{1}$H NMR (800 MHz) spectrum of **2** in $CDCl_3$. | 25 |
| **Fig. 16.** HSQC (800 MHz) spectrum of **2** in $CDCl_3$. | 26 |
| **Fig. 17.** DQF-COSY (800 MHz) spectrum of **2** in $CDCl_3$. | 26 |
| **Fig. 18.** HMBC (800 MHz) spectrum of **2** in $CDCl_3$. | 27 |
| **Fig. 19.** NOESY (800 MHz) spectrum of **2** in $CDCl_3$. | 27 |
| **Fig. 20.** IR spectrum of **2**. | 28 |
| **Fig. 21.** Multiple sequence alignment of KS domains. | 29 |
| **Fig. 22.** Multiple sequence alignment of AT domains. | 30 |
| **Fig. 23.** Multiple sequence alignment of KR domains. | 31 |
| **Fig. 24.** Multiple sequence alignment of DH domains. | 32 |
| **Fig. 25.** Multiple sequence alignment of ER domains. | 33 |
| **Fig. 26.** Sequence alignment of AliB, AliK, AliL and other cytochrome P450s. | 34 |
| **Fig. 27.** The genome editing of tailoring enzymes in alligamycin A biosynthesis. | 35 |
| **Fig. 28.** Sequence similarity network of AliJ. | 36 |
| **Fig. 29.** Multiple sequence alignment of AliH and other crotonyl-CoA carboxylase/reductases. | 37 |
| **Fig. 30.** The COGs functional classification of differential proteins in *A. niger*. | 38 |
| **Fig. 31.** The GO annotation of differential proteins in *A. niger*. | 39 |
| **Fig. 32.** The KEGG pathway enrichment analysis of differential proteins in *A. niger*. | 40 |

| | |
|---|---|
| **Fig. 33.** The abundance of responsive protein of known antifungal drugs in alligamycin A treatment. | 40 |
| **References** | 41 |

**Tab. 1.** All detected BGCs belonging to the GCF_00315 in BiG-FAM database.

| **Dataset** | **BGC** | **Distance to model** | **Length (kb)** | **Taxon** |
|---|---|---|---|---|
| MIBiG | BGC0001670.1 | 640 | 88.86 | *Streptomyces cinnamonensis* |
| MIBiG | BGC0000058.1 | 712 | 74.32 | *Streptomyces graminofaciens* |
| Isolated bacterial draft | GCF_001941325.1/NZ_LYDV01000578.region001 | 742 | 28.08 | *Streptomyces acidiscabies* |
| Isolated bacterial draft | GCF_001941265.1/NZ_LYDT01000970.region001 | 760 | 19.82 | *Bacteria* (Kingdom) |
| MIBiG | BGC0000114.1 | 760 | 95.7 | *Streptomyces violaceusniger* |
| Isolated bacterial draft | GCF_010548575.1/c00095_NZ_JAAG...region001 | 794 | 44.15 | *Streptomyces* (Genus) |
| Isolated bacterial draft | GCF_001949645.1/NZ_BCML01000004.region001 | 819 | 63.06 | *Streptomyces acidiscabies* |
| Isolated bacterial draft | GCF_001485105.1/NZ_BCMK01000018.region001 | 826 | 63.14 | *Streptomyces acidiscabies* |
| MIBiG | BGC0000109.1 | 835 | 80.36 | *Streptomyces cyaneogriseus* |
| MIBiG | BGC0000100.1 | 844 | 103.45 | *Streptomyces cinnamonensis* |

**Tab. 2.** The biosynthetic gene clusters in *S. iranensis* based on genome resequencing.

| Region | Type | Position | | Most similar known cluster | Similarity |
|---|---|---|---|---|---|
| Region 1 | lassopeptide | 47,826 | 67,848 | SSV-2083 | 18% |
| Region 2 | NRPS-like, T1PKS, NRPS | 89,243 | 264,517 | meridamycin | 10% |
| Region 3 | terpene, NRPS | 310,755 | 369,526 | carotenoid | 63% |
| Region 4 | NRPS | 584,615 | 633,798 | coelichelin | 100% |
| Region 5 | butyrolactone | 920,071 | 929,026 | cyphomycin | 9% |
| Region 6 | phosphonate, acyl_amino_acids, butyrolactone, NRPS-like, T1PKS, hserlactone | 983,576 | 1,129,298 | cyphomycin | 5% |
| Region 7 | T1PKS | 1,216,733 | 1,353,478 | azalomycin F3a | 100% |
| Region 8 | T1PKS | 1,525,866 | 1,708,371 | nigericin | 70% |
| Region 9 | T1PKS | 1,812,276 | 1,890,013 | efomycin K/F | 95% |
| Region 10 | redox-cofactor | 1,914,682 | 1,936,746 | | |
| Region 11 | hserlactone | 2,054,025 | 2,074,780 | heronamides | 8% |
| Region 12 | butyrolactone | 2,085,494 | 2,096,426 | | |
| Region 13 | T1PKS, NRPS | 2,274,879 | 2,327,099 | meilingmycin | 4% |
| Region 14 | NRPS, T3PKS, other | 2,383,216 | 2,482,446 | feglymycin | 78% |
| Region 15 | NRPS, T1PKS | 2,535,309 | 2,739,409 | kitacinnamycin | 16% |
| Region 16 | terpene | 2,985,464 | 3,007,479 | hopene | 76% |
| Region 17 | T1PKS | 3,153,626 | 3,235,531 | bafilomycin B1 | 55% |
| Region 18 | T2PKS | 3,304,807 | 3,374,688 | spore pigment | 83% |
| Region 19 | T1PKS | 3,499,162 | 3,559,357 | notonesomycin A | 11% |
| Region 20 | RiPP-like | 3,612,469 | 3,622,637 | | |
| Region 21 | siderophore | 3,792,665 | 3,803,134 | | |
| Region 22 | T2PKS | 4,140,701 | 4,213,204 | isoindolinomycin | 61% |
| Region 23 | NRPS-like | 4,442,739 | 4,484,596 | echosides | 100% |
| Region 24 | siderophore | 5,229,788 | 5,240,857 | legonoxamines/ desferrioxamin B | 83% |
| Region 25 | terpene | 6,441,503 | 6,461,832 | TVA-YJ-2 | 9% |
| Region 26 | ladderane, arylpolyene, NRPS | 6,705,172 | 6,807,080 | kitacinnamycins | 50% |
| Region 27 | NRPS | 7,345,535 | 7,387,933 | ochronotic pigment | 75% |
| Region 28 | ladderane | 7,449,836 | 7,489,866 | cinnapeptin | 53% |
| Region 29 | RRE-containing | 7,510,172 | 7,530,691 | granaticin | 10% |
| Region 30 | T1PKS | 7,664,407 | 7,848,116 | desulfoclethramycin /clethramycin | 59% |
| Region 31 | terpene | 8,396,931 | 8,416,950 | | |
| Region 32 | ectoine | 9,062,415 | 9,072,819 | ectoine | 100% |
| Region 33 | siderophore | 9,237,702 | 9,251,453 | peucechelin | 20% |
| Region 34 | terpene | 9,301,372 | 9,319,391 | BE-43547s | 25% |
| Region 35 | nucleoside | 9,535,814 | 9,572,869 | huimycin | 100% |
| Region 36 | nucleoside, NRPS, T1PKS, NRPS-like | 9,677,504 | 9,821,457 | rapamycin | 82% |
| Region 37 | PKS-like | 9,909,290 | 9,950,318 | rustmicin | 33% |

| | | | | | |
|---|---|---|---|---|---|
| Region 38 | terpene | 10,000,500 | 10,020,321 | 2-methylisoborneol | 100% |
| Region 39 | terpene | 10,464,893 | 10,484,016 | pristinol | 100% |
| Region 40 | T1PKS, ladderane,arylpolyene | 10,578,295 | 10,629,355 | cinnapeptin | 46% |
| Region 41 | T1PKS, NRPS-like | 10,835,982 | 10,918,034 | hygrocins | 83% |
| Region 42 | lanthipeptide-class-ii | 11,011,708 | 11,035,079 | reveromycin A | 9% |
| Region 43 | hglE-KS, T1PKS, RiPP-like | 11,062,970 | 11,124,110 | Hexacosalactone A | 68% |
| Region 44 | T1PKS, NRPS | 11,127,758 | 11,269,230 | neomediomycin B | 32% |
| Region 45 | terpene | 11,320,894 | 11,341,820 | brasilicardin A | 30% |
| Region 46 | terpene | 11,470,415 | 11,491,323 | | |
| Region 47 | T1PKS, NRPS-like | 11,577,719 | 11,624,306 | niphimycins C-E | 12% |
| Region 48 | betalactone | 11,818,784 | 11,847,595 | Sch-47554 / Sch-47555 | 7% |
| Region 49 | NRPS, NRPS-like, T1PKS | 12,040,448 | 12,123,871 | notonesomycin A | 3% |
| Region 50 | lassopeptide | 12,143,153 | 12,165,683 | SSV-2083 | 18% |

**Tab. 3.** The function annotation of biosynthetic genes of *ali* BGC.

| Gene | Locus tag[a] | Size[b] | Proposed functions | SI/ID[c] | Protein homologue and origin |
|---|---|---|---|---|---|
| *aliA* | RY074_00730 | 5764 | Type I polyketide synthase | 70/77 | WP_069850026.1, *Actinoalloteichus hymeniacidonis* |
| *aliB* | RY074_00725 | 416 | Cytochrome P450 | 68/79 | WP_028677909.1, *Salinispora arenicola* |
| *aliC* | RY074_00720 | 4443 | Type I polyketide synthase | 54/65 | WP_098246318.1, *Streptomyces formicae* |
| *aliD* | RY074_00715 | 8121 | Type I polyketide synthase | 55/67 | WP_100583907.1, *Streptomyces* sp. CB02120-2 |
| *aliE* | RY074_00710 | 5720 | Type I polyketide synthase | 49/60 | BAE93731.1, *Streptomyces* sp. NRRL 11266 |
| *aliF* | RY074_00705 | 1969 | Type I polyketide synthase | 56/66 | WP_116210512.1, *Streptomyces olivoreticuli* |
| *aliG* | RY074_00700 | 269 | Methyltransferase domain-containing protein | 65/75 | WP_198270846.1, *Streptomyces sabulosicollis* |
| *aliH* | RY074_00695 | 411 | Crotonyl-CoA carboxylase/reductase | 74/85 | WP_042181611.1, *Kibdelosporangium* sp. MJ126-NF4 |
| *aliI* | RY074_00690 | 654 | Hypothetical protein | 42/54 | WP_089101481.1, *Streptomyces hyaluromycini* |
| *aliJ* | RY074_00685 | 338 | UbiD family decarboxylase | 60/75 | WP_106196453.1, *Umezawaea tangerina* |
| *aliK* | RY074_00680 | 394 | Cytochrome P450 | 58/72 | MBR7672783.1, *Streptomyces daliensis* |
| *aliL* | RY074_00675 | 403 | Cytochrome P450 | 52/68 | WP_227725975.1, *Streptomyces* sp. ET3-23 |

[a], The gene locus tags as in NCBI GenBank accession CP136563.1; [b], The size of amino acids; [c], similarity-identity ratio; *, located in the inverted terminal repeat region.

**Tab. 4.** $^{1}H$ (800 MHz) and $^{13}C$ (200 MHz) NMR Data for alligamycin A (**1**) and alligamycin B (**2**) in $CDCl_3$.

| pos. | alligamycin A | | | | alligamycin B | | | |
|---|---|---|---|---|---|---|---|---|
| | $\delta_C$, type | $\delta_H$, (*J* in Hz) | COSY | HMBC | $\delta_C$, type | $\delta_H$, (*J* in Hz) | COSY | HMBC |
| 1 | 168.6, C | - | | | 168.3, C | - | | |
| 2 | 128.5, CH | 6.19 (d, 15.4) | 3 | 1, 4, 5 | 125.9, CH | 6.11 (d, 15.7) | 3 | 1(w), 4 |
| 3 | 137.8, CH | 7.91 (dd, 15.4, 11.8) | 2, 4 | 1, 2, 5 | 141.0, CH | 7.94 (dd, 15.7,11.7) | 2, 4 | 1, 5(w) |
| 4 | 129.1, CH | 6.47 (d, 10.7) | 3 | 2, 6, 40 | 133.6, CH | 6.74 (d, 11.6) | 3 | 2, 6, 40 |
| 5 | 143.2, C | | | | 144.2, C | - | | |
| 6 | 79.4, CH | 4.88 (d, 8.9) | 7 | 4, 5, 8, 40, 41 | 75.8, CH | 4.35 (d, 8.4) | 7 | 4, 5, 7, 8, 40 |
| 7 | 41.5, CH | 1.93 (overlapping) | 6, 8, 41 | 5, 6, 41 | 40.7, CH | 1.98 (overlapping) | 8 | 6 |
| 8 | 76.5, CH | 3.37 (overlapping) | 7, 9 | 41, 47 | 79.6, CH | 3.45 (m) | 7, 9 | |
| 9 | 32.0, $CH_2$ | 1.51 (d, 4.7),<br>1.46 (d, 4.10) | 8, 10 | 8, 10 | 33.6, $CH_2$ | 1.58 (m) | 9, 10 | |
| 10 | 71.2, CH | 5.15 (dq, 3.4, 2.1) | 9, 11 | 8, 9, 42, 46 | 72.0, CH | 5.14 (dq, 3.4, 2.1) | 9, 11 | |
| 11 | 48.4, CH | 3.03 (m) | 10, 42 | 10, 12, 42 | 48.7, CH | 2.97 (m) | 10 | 10, 12(w) |
| 12 | 207.3, C | | | | 207.3, C | - | | |
| 13 | 47.8, $CH_2$ | 2.81 (dd, 12.4, 2.9),<br>2.50 (dd, 12.4, 10.1) | | 11, 12, 14, 15 | 47.9, $CH_2$ | 2.73 (dd, 13.2, 3.9),<br>2.53 (dd, 13.2, 9.2) | 14 | 12, 14, 15 |
| 14 | 60.7, CH | 4.12 (overlapping) | 13, 15 | | 60.7, CH | 4.13 (overlapping) | 13, 15 | |
| 15 | 38.2, $CH_2$ | 1.93 (overlapping),<br>1.51 (overlapping) | 14, 16 | 14 | 37.1, $CH_2$ | 1.93 (overlapping),<br>1.51 (overlapping) | 14, 16 | 14 |
| 16 | 65.0, CH | 4.15 (overlapping) | 15, 17 | | 64.9, CH | 4.14 (overlapping) | 15, 17 | |
| 17 | 37.0, $CH_2$ | 1.97 (d, 14.1),<br>1.45 (dd, 14.2, 3.5) | 16 | 15, 16, 18 | 37.0, $CH_2$ | 1.97 (d, 14.1),<br>1.45 (dd, 14.2, 3.5) | 16 | 18(w) |
| 18 | 101.0, C | - | | | 101.0, C | - | | |
| 19 | 34.8, CH | 1.66 (m) | 20, 43 | 18, 20 | 34.8, CH | 1.66 (m) | 43 | 18(w), 20 |
| 20 | 25.4, $CH_2$ | 2.10 (overlapping) | 19, 21 | 19, 21 | 25.4, $CH_2$ | 2.10 (overlapping) | 21 | |

| | | | | | | | | |
|---|---|---|---|---|---|---|---|---|
| 21 | 25.4, $CH_2$ | 1.40 (overlapping), 1.28 (overlapping) | 20 | 22 | 25.4, $CH_2$ | 1.40 (overlapping), 1.25 (overlapping) | 20, 22 | |
| 22 | 72.0, CH | 3.23 (dt, 11,5, 9.6) | 21, 23 | 23 | 72.0, CH | 3.23 (dt, 11,5, 9.6) | 21, 23 | |
| 23 | 33.6, $CH_2$ | 1.58 (m), 1.26 (overlapping) | 22, 24 | 22 | 33.6, $CH_2$ | 1.58 (m), 1.26 (overlapping) | 22, 24 | 22 |
| 24 | 33.9, $CH_2$ | 1.14 (m) | 23, 25 | 23, 25, 44 | 34.0, $CH_2$ | 1.15 (m) | 23, 25 | 22, 25, 44 |
| 25 | 30.0, CH | 2.15 (overlapping) | 24, 26, 44 | | 30.0, CH | 2.15 (overlapping) | 24, 26 | |
| 26 | 51.8, $CH_2$ | 2.61 (dd, 16.3, 10.7), 2.24 (dd, 16.3, 7.8) | 25 | 25, 27, 28, 44 | 51.8, $CH_2$ | 2.61 (dd, 16.0, 10.5), 2.24 (dd, 16.3, 7.8) | 25 | 24, 25, 27, 44 |
| 27 | 201.3, C | - | | | 201.6, C | - | | |
| 28 | 123.0, CH | 6.48 (s) | | 27, 29, 30, 45 | 123.2, CH | 6.52 (s) | | 29, 30, 45 |
| 29 | 156.4, C | - | | | 156.4, C | - | | |
| 30 | 75.8, CH | 4.14 (overlapping) | 31 | 28, 29, 31, 32, 45, 46 | 75.5, CH | 4.14 (overlapping) | 31 | 29, 31, 46 |
| 31 | 47.2, CH | 2.83 (dd, 8.8, 2.4) | 30, 32 | 32, 33, 46 | 47.2, CH | 2.83 (dt, 8.8, 2.4) | 30, 32 | 32, 46 |
| 32 | 30.7, $CH_2$ | 1.93 (overlapping), 1.81 (m) | 31, 33 | 31, 33, 34, 46 | 30.7, $CH_2$ | 1.93 (overlapping), 1.81 (m) | 31 | 33(w), 34(w), 46 |
| 33 | 27.4, $CH_2$ | 1.48 (overlapping), 1.34 (overlapping) | 32 | | 27.4, $CH_2$ | 1.48 (overlapping), 1.34 (overlapping) | | |
| 34 | 29.2, $CH_2$ | 1.38 (overlapping) | | | 29.2, $CH_2$ | 1.38 (overlapping) | | |
| 35 | 28.8, $CH_2$ | 1.33 (overlapping) | 36 | | 28.8, $CH_2$ | 1.33 (overlapping) | 36 | |
| 36 | 23.6, $CH_2$ | 1.58 (m) | 35, 37 | 35, 37, 38 | 23.6, $CH_2$ | 1.58 (m) | 35 | 35, 37, 38 |
| 37 | 43.6, $CH_2$ | 2.46 (t,7.4) | 36 | 35, 36, 38 | 43.6, $CH_2$ | 2.46 (t, 7.4) | 36 | 35, 36, 38 |
| 38 | 209.6, C | - | | | 209.6, C | - | | |
| 39 | 29.9, $CH_3$ | 2.15 (s) | | 37, 38 | 30.0, $CH_3$ | 2.13 (s) | | 37, 38 |
| 40 | 161.8, C | - | | | 166.9, C | | | |
| 41 | 8.4, $CH_3$ | 0.94 (d, 7.2) | 7 | 6, 7, 8 | 11.6, $CH_3$ | 0.77 (d, 7.2) | 7 | 6, 7, 8 |
| 42 | 9.2, $CH_3$ | 0.95 (d,6.8) | 11 | 10, 11, 12 | 9.6, $CH_3$ | 0.96 (d,7.2) | 11 | 10, 11, 69 |

| | | | | | | | | |
|---|---|---|---|---|---|---|---|---|
| 43 | 14.3, $CH_3$ | 0.98 (d, 7.2) | 19 | 18, 19, 20 | 14.3, $CH_3$ | 0.87 (d, 7.2) | 19 | 18, 19, 20 |
| 44 | 19.5, $CH_3$ | 0.93 (d, 6.8) | 25 | 25, 26 | 19.5, $CH_3$ | 0.92 (d, 6.8) | 25 | 24, 25, 26 |
| 45 | 17.4, $CH_3$ | 2.10 (s) | | 27, 28, 29, 30 | 17.3, $CH_3$ | 2.07 (s) | | 27(w), 28, 29, 30 |
| 46 | 174.9, C | - | | | 175.0, C | - | | |
| 8-OMe | 59.6, $CH_3$ | 3.37 (s) | | 8 | 58.7, $CH_3$ | 3.36 (s) | | 8 |
| 40-OMe | | | | | 52.1, $CH_3$ | 3.85 (s) | | 40 |

**Tab. 5.** Alignment of conserved motifs in the active site of AT domains.

| domain name | specificity | 198 | 199 | 200 | 201 |
|---|---|---|---|---|---|
| ave_AT5 | Malonyl-CoA | H | A | F | H |
| nid_AT3 | Malonyl-CoA | H | A | F | H |
| epo_AT2 | Malonyl-CoA | H | A | F | H |
| amp_AT18 | Malonyl-CoA | H | A | F | H |
| rif_AT2 | Malonyl-CoA | H | A | F | H |
| ali_AT1 | Malonyl-CoA | H | A | F | H |
| ali_AT2 | Malonyl-CoA | H | A | F | H |
| ali_AT5 | Malonyl-CoA | H | A | F | H |
| ali_AT7 | Malonyl-CoA | H | A | F | H |
| ali_AT8 | Malonyl-CoA | H | A | F | H |
| ali_AT9 | Malonyl-CoA | H | A | F | H |
| ali_AT11 | Malonyl-CoA | H | A | F | H |
| ali_AT12 | Malonyl-CoA | H | A | F | H |
| ali_AT14 | Malonyl-CoA | H | A | F | H |
| ave_AT1 | Methylmalony-CoA | Y | A | S | H |
| ery_AT4 | Methylmalony-CoA | Y | A | S | H |
| amp_AT2 | Methylmalony-CoA | Y | A | S | H |
| rif_AT7 | Methylmalony-CoA | Y | A | S | H |
| ali_AT3 | Methylmalony-CoA | Y | A | S | H |
| ali_AT4 | Methylmalony-CoA | Y | A | S | H |
| ali_AT5 | Methylmalony-CoA | Y | A | S | H |
| ali_AT10 | Methylmalony-CoA | Y | A | S | H |
| ali_AT13 | Methylmalony-CoA | Y | A | S | H |
| ali_AT15 | Methylmalony-CoA | Y | A | S | H |
| nid_AT5 | Ethylmalonyl-CoA | T | A | G | H |
| tyl_AT5 | Ethylmalonyl-CoA | T | A | G | H |
| ali_AT16 | Malonyl-CoA | I | A | S | H |

Abbreviations: ali, alligamycin; ave, avermectin; nid, niddamycin; epo, epothilone; amp, amphotericin; rif, rifamycin; ery, erythromycin; tyl, tylactone; meg, megalomicin; pta, pteridic acids.

**Tab. 6.** Antifungal assays of alligamycin A (**1**).

| **Strain number** | **Species** | **Section** | **Amphotericin B MIC in mg/L** | **Itraconazole MIC in mg/L** | **Voriconazole MIC in mg/L** | **Alligamycin A (1) MIC (MEC) in mg/L** |
|---|---|---|---|---|---|---|
| NRZ-2019-445 | *Aspergillus flavus* | Flavi | 4 | 0.5 | 0.5 | >8 (0.125) |
| NRZ-2023-1167 | *Aspergillus flavus* | Flavi | 8 | 0.5 | 1 | >8 (0.5) |
| SF006305 | *Aspergillus parasiticus* | Flavi | 2 | 0.5 | 0.5 | >8 (0.25) |
| NRZ-2021-0631 | *Aspergillus lentulus* | Fumigati | 8 | 1 | 2 | >8 (>8) |
| NRZ-2022-0606 | *Aspergillus fumigatus* (Azole-R) | Fumigati | 1 | 8 | 2 | >8 (>8) |
| ATCC204305 | *Aspergillus fumigatus* (Azole-S) | Fumigati | 1 | 1 | 0.5 | >8 (>8) |
| NRZ-2020-060 | *Aspergillus udagawae* | Fumigati | 4 | 8 | 8 | >8 (8) |
| NRZ-2023-0949 | *Aspergillus latus* | Nidulantes | 0.25 | 0.5 | 0.25 | 4 (0.5) |
| NRZ-2023-0770 | *Aspergillus nidulans* | Nidulantes | 4 | 0.25 | 0.125 | 4 (2) |
| NRZ-2023-0889 | *Aspergillus spinulosporus* | Nidulantes | 0.5 | 0.5 | 0.25 | 8 (0.25) |
| SF011948 | *Aspergillus brasiliensis* | Nigri | 0.5 | 2 | 2 | 0.5 (0.25) |
| NRZ-2023-0685 | *Aspergillus luchuensis* | Nigri | 0.25 | n.a. | 1 | 0.5 (0.25) |
| NRZ-2022-0820 | *Aspergillus niger* | Nigri | 0.5 | n.a. | 1 | 4 (0.5) |
| NRZ-2022-0863 | *Aspergillus tubingensis* | Nigri | 0.5 | 4 | 1 | >8 (0.25) |
| NRZ-2023-0826 | *Aspergillus welwitschiae* | Nigri | 0.25 | n.a. | 0.5 | 1 (0.25) |
| NRZ-2021-0572 | *Aspergillus citrinoterreus* | Terrei | 4 | 0.5 | 0.5 | 0.5 (0.25) |
| SF013938 | *Aspergillus neoafricanus* | Terrei | 2 | 0.5 | 0.5 | 0.5 (0.25) |
| NRZ-2019-093 | *Aspergillus terreus* | Terrei | 8 | 0.5 | 1 | 0.125 (0.125) |
| NRZ-2019-116 | *Aspergillus calidoustus* | Usti | 16 | 8 | 4 | 0.5 (0.25) |
| SF006408 | *Aspergillus ustus* | Usti | 1 | >8 | >8 | 0.125 (0.06) |
| NRZ-2023-0832 | *Candida albicans* (Echino-R) | | 0.125 | ≤0.016 | ≤0.016 | >8 |
| NRZ-2023-0868 | *Candida albicans* (Echino-S) | | 0.06 | 0.03 | ≤0.016 | >8 |
| NRZ-2023-0919 | *Candida auris* | | 1 | 0.06 | 0.125 | >8 |

| | | | | | |
|---|---|---|---|---|---|
| NRZ-2023-0642 | *Candida glabrata* (Echino-R) | 0.5 | 1 | 0.25 | >8 |
| NRZ-2023-0711A | *Candida glabrata* (Echino-S) | 0.5 | 1 | 0.25 | >8 |
| NRZ-2023-0854 | *Candida parapsilosis* (Azol-R) | 0.25 | 0.125 | 0.25 | >8 |
| NRZ-2023-0829 | *Candida parapsilosis* (Azol-S) | 0.25 | 0.06 | ≤0.016 | 8 |
| NRZ-2020-713 | *Fusarium annulatum* | 2 | >8 | 8 | >8 (2) |
| NRZ-2020-259 | *Fusarium musae* | 4 | >8 | 4 | >8 (4) |
| NRZ-2020-699 | *Lomentospora prolificans* | >8 | >8 | >8 | >8 (>8) |
| NRZ-2023-0726 | *Mucor circinelloides* | 0.06 | 2 | >8 | >8 (>8) |
| NRZ-2022-0133 | *Purpureocilium lilacinum* | >16 | >8 | 0.25 | >8 (>8) |
| NRZ-2023-0992 | *Rhizopus arrhizus* | 0.125 | 1 | 4 | >8 (>8) |
| NRZ-2023-0599 | *Talaromyces columbinus* | 2 | >8 | 8 | >8 (0.5) |
| NRZ-2021-0639 | *Talaromyces kabodanensis* | 0.25 | >8 | 8 | 2 (0.5) |
| NRZ-2016-235 | *Talaromyces marneffei* | 0.06 | ≤0.016 | ≤0.016 | 0.125 (0.125) |
| NRZ-2021-0646 | *Talaromyces purpureogenes* | 1 | >8 | 8 | 0.5 (0.25) |
| NRZ-2022-0441 | *Trichoderma longibrachiatum* | 2 | >8 | 1 | >8 (>8) |

MIC: minimum inhibitory concentration.

**Tab. 7.** The strains and plasmids used in this study.

| Strains | Description | Source/[Ref] |
|---|---|---|
| One Shot™ Mach1™ T1 Phage-Resistant Chemically Competent *E. coli* | For routine plasmids maintenance and cloning | Thermo Fisher Scientific |
| *E. coli* ET12567/pUZ8002 | For conjugating plasmids into *Streptomyces* | [1] |
| *S. iranensis* | Wild-type strain | DSMZ |
| *S. iranensis*/Δ*aliA* | Knock-out of *aliA* in *S. iranensis* | In this work |
| *S. iranensis*/Δ*aliB* | Inactivation of *aliB* in *S. iranensis* | In this work |
| *S. iranensis*/Δ*aliG* | Inactivation of *aliG* in *S. iranensis* | In this work |
| *S. iranensis*/Δ*aliH* | Inactivation of *aliH* in *S. iranensis* | In this work |
| *S. iranensis*/Δ*aliI* | Inactivation of *aliI* in *S. iranensis* | In this work |
| *S. iranensis*/Δ*aliJ* | Knock-out of *aliJ* in *S. iranensis* | In this work |
| *S. iranensis*/Δ*aliK* | Inactivation of *aliK* in *S. iranensis* | In this work |
| *S. iranensis*/Δ*aliL* | Inactivation of *aliL* in *S. iranensis* | In this work |
| *S. iranensis*/Δ*aliH::aliH* | Complementation strain of Δ*aliH* mutant | In this work |
| **Plasmids** | | |
| pCRISPR-Cas9 | Gene knockout/in for actinomycetes | [2] |
| pCRISPR-cBEST | For C to T base editing | [3] |
| pGM1190 | Plasmid for gene complementation | [4] |
| pCRISPR-Cas9/Δ*aliA* | Modified plasmid for knock-out of *aliA* | In this work |
| pCRISPR-Cas9/Δ*aliJ* | Modified plasmid for knock-out of *aliJ* | In this work |
| pCRISPR-cBEST/Δ*aliB* | Modified plasmid for inactivation of *aliB* | In this work |
| pCRISPR-cBEST/Δ*aliG* | Modified plasmid for inactivation of *aliG* | In this work |
| pCRISPR-cBEST/Δ*aliH* | Modified plasmid for inactivation of *aliH* | In this work |
| pCRISPR-cBEST/Δ*aliI* | Modified plasmid for inactivation of *aliI* | In this work |
| pCRISPR-cBEST/Δ*aliK* | Modified plasmid for inactivation of *aliK* | In this work |
| pCRISPR-cBEST/Δ*aliL* | Modified plasmid for inactivation of *aliL* | In this work |
| pGM1190-*aliH* | Modified plasmid for complementation | In this work |

**Tab. 8.** The primers used in this study.

| **Primer name** | **Sequence (5' → 3')** | **Description** |
|---|---|---|
| Del-*aliA*-sgRNA | CCGGTTGGTAGGATCGACGGgaccagtggcctcc gcgtaGTTTTAGAGCTAGAAATAGC | Knock-out of *aliA,* the base marked in red is sgRNA sequence |
| Del-*aliA*-uparm-F | tcgtcgaaggcactagaaggatctggttgcagtcgtcgta | Forward primer for amplification of upstream homologous flank of *aliA* |
| Del-*aliA*-uparm-R | actcacctgttccagtgatccacttgcacgaactgaccag | Reverse primer for amplification of upstream homologous flank of *aliA* |
| Del-*aliA*-downarm-F | ctggtcagttcgtgcaagtggatcactggaacaggtgagt | Forward primer for amplification of downstream homologous flank of *aliA* |
| Del-*aliA*-downarm-R | ggtcgatccccgcatatagggtacatgatcgacggacaag | Reverse primer for amplification of downstream homologous flank of *aliA* |
| ID-*aliA*-F1 | atgaagaagatcgagctgatg | Forward primer for screening *aliA* mutants |
| ID-*aliA*-R1 | ctcaacgcgttccagatc | Reverse primer for screening *aliA* mutants |
| ID-*aliA*-F2 | cttcgaggagctcgtcgtac | Forward primer for screening *aliA* mutants |
| ID-*aliA*-R2 | catctgttaacggcacacc | Reverse primer for screening *aliA* mutants |
| Del-*aliJ*-sgRNA | CCGGTTGGTAGGATCGACGGtcgagaagtccccgat catgGTTTTAGAGCTAGAAATAGC | Knock-out of *aliJ*, the base marked in red is sgRNA sequence |
| Del-*aliJ*-uparm-F | tcgtcgaaggcactagaagggcgcaaactggtcaacatggc | Forward primer for amplification of upstream homologous flank of *aliJ* |
| Del-*aliJ*-uparm-R | gagagatcgacggcgacatggagcatccacgtgctgaccgacaacc | Reverse primer for amplification of upstream homologous flank of *aliJ* |
| Del-*aliJ*-downarm-F | ggttgtcggtcagcacgtggatgctccatgtcgccgtcgatctctc | Forward primer for amplification of downstream homologous flank of *aliJ* |
| Del-*aliJ*-downarm-R | ggtcgatccccgcatataggctgaccgactacctgagcacgg | Reverse primer for amplification of downstream homologous flank of *aliJ* |
| ID-*aliJ*-F1 | gacctcgaagacggactcg | Forward primer for screening *aliJ* mutants |
| ID-*aliJ*-R1 | gaagagtccgatcaggaag | Reverse primer for screening *aliJ* mutants |
| ID-*aliJ*-F2 | ctccctgatcttcaacaagctc | Forward primer for screening *aliJ* mutants |
| ID-*aliJ*-R2 | ttcccgctggccttcaac | Reverse primer for screening *aliJ* mutants |
| Inact-*aliB* | CCGGTTGGTAGGATCGACGGcaaccagatcctgcgc cacaGTTTTAGAGCTAGAAATAGC | Inactivation of *aliB*, the base marked in red is sgRNA sequence |
| Inact-*aliG* | CCGGTTGGTAGGATCGACGGcggtccagtagccga ggtgcGTTTTAGAGCTAGAAATAGC | Inactivation of *aliG*, the base marked in red is sgRNA sequence |
| Inact-*aliH* | CCGGTTGGTAGGATCGACGGgctcccagccacacaa ctgcGTTTTAGAGCTAGAAATAGC | Inactivation of *aliH*, the base marked in red is sgRNA sequence |
| Inact-*aliI* | CCGGTTGGTAGGATCGACGGgaaccagaccttcagc gtacGTTTTAGAGCTAGAAATAGC | Inactivation of *aliI*, the base marked in red is sgRNA sequence |
| Inact-*aliK* | CCGGTTGGTAGGATCGACGGctgatccaggcccgtg acgaGTTTTAGAGCTAGAAATAGC | Inactivation of *aliK*, the base marked in red is sgRNA sequence |
| Inact-*aliL* | CCGGTTGGTAGGATCGACGGaactcccagagtccgt agaaGTTTTAGAGCTAGAAATAGC | Inactivation of *aliL*, the base marked in red is sgRNA sequence |
| ID-*aliB*-F | cagcatcgtgcggaagatc | Forward primer for screening *aliB* mutants |

| | | |
|---|---|---|
| ID-*aliB*-R | ctgctcgctctgctgga | Reverse primer for screening *aliB* mutants |
| ID-*aliG*-F | cttgtctagttcctggatcc | Forward primer for screening *aliG* mutants |
| ID-*aliG*-R | gtcttcgactatcccacg | Reverse primer for screening *aliG* mutants |
| ID-*aliH*-F | gacgatctggatggccatc | Forward primer for screening *aliH* mutants |
| ID-*aliH*-R | catatgacacggaccgtgtac | Reverse primer for screening *aliH* mutants |
| ID-*aliI*-F | actcccttgacgttcttcttg | Forward primer for screening *aliI* mutants |
| ID-*aliI*-R | ctacgtctggctcatgaacac | Reverse primer for screening *aliI* mutants |
| ID-*aliK*-F | tggtcaagaactcgccgaac | Forward primer for screening *aliK* mutants |
| ID-*aliK*-R | aagcggagtacctcctcgac | Reverse primer for screening *aliK* mutants |
| ID-*aliL*-F | catccaggagatcgtcgac | Forward primer for screening *aliL* mutants |
| ID-*aliL*-R | cgtgccgttcaccaataac | Reverse primer for screening *aliL* mutants |
| pGM1190-*aliH*-F | agaagggagcggacatatgaatgacacggaccgtgtacga | Forward primer for amplification of *aliH* |
| pGM1190-*aliH*-R | acaaaactttagatctggggctagttcctggatccgggtt | Reverse primer for amplification of *aliH* |
| ID-pGM1190-*aliH*-F | gaggtcattactggaccgg | Forward primer for verification of *aliH* complementation |
| ID-pGM1190-*aliH*-R | cactccgctgaaactgttg | Reverse primer for verification of *aliH* complementation |

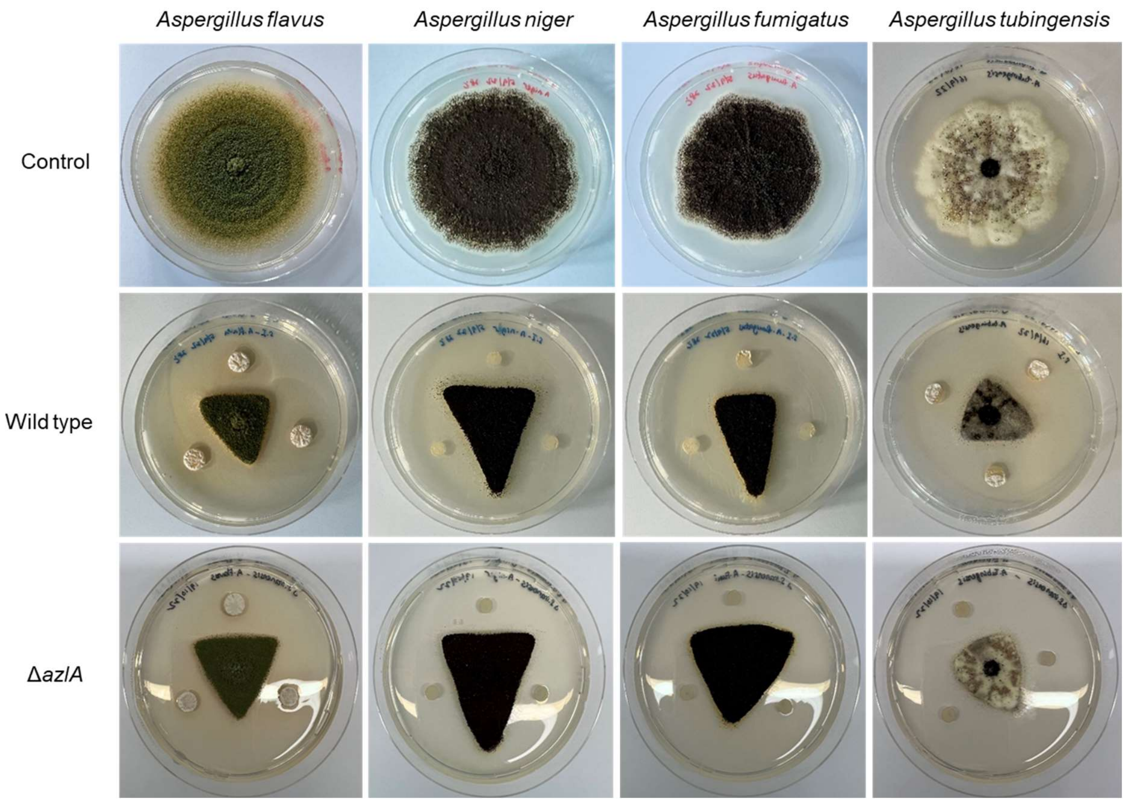


**Fig. 1.** Co-culture of the wild type or azalomycins-deficient mutant Δ*azlA* of *S. iranensis* (individual colonies scattered in three corners) with different *Aspergillus* strains (middle) on PDB agar medium.

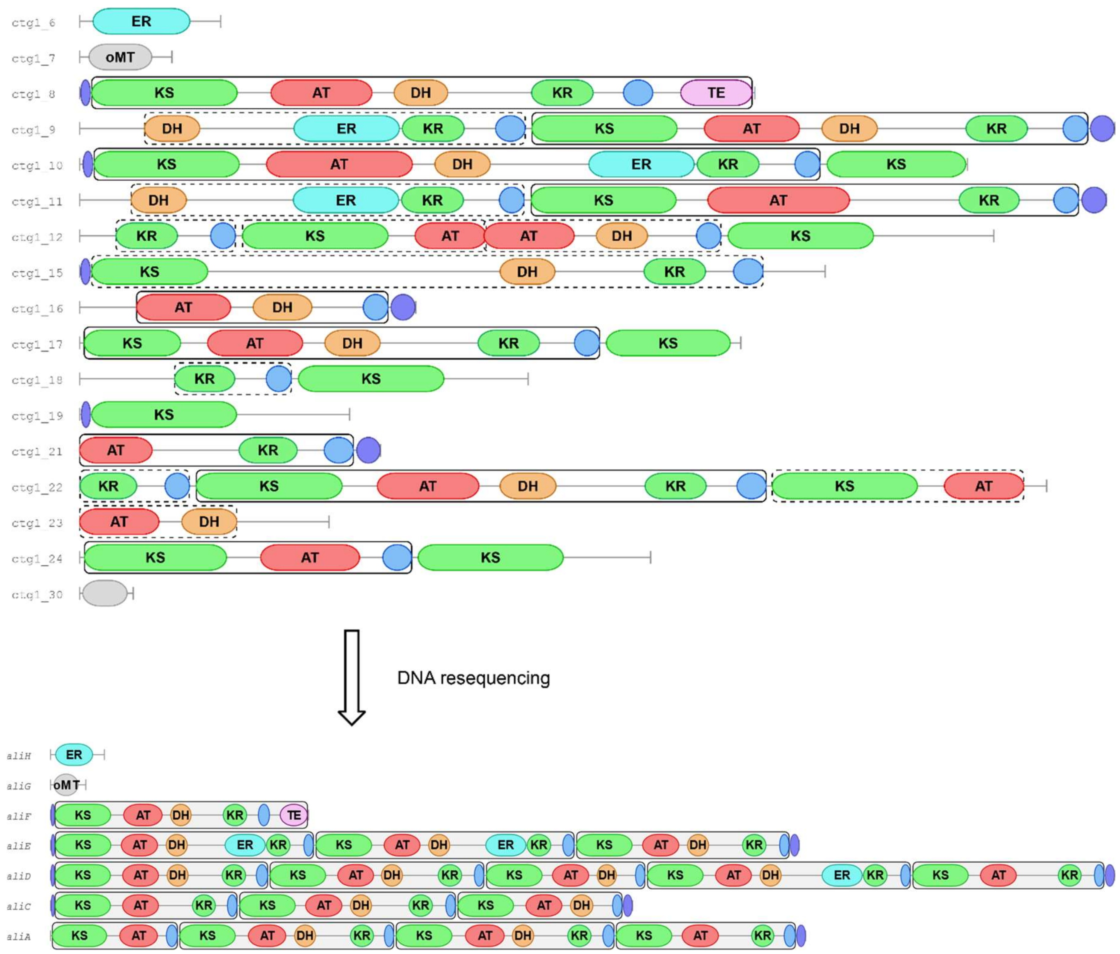


**Fig. 2.** Domain annotation of *ali* BGC in *S. iranensis* before and after whole-genome resequencing.

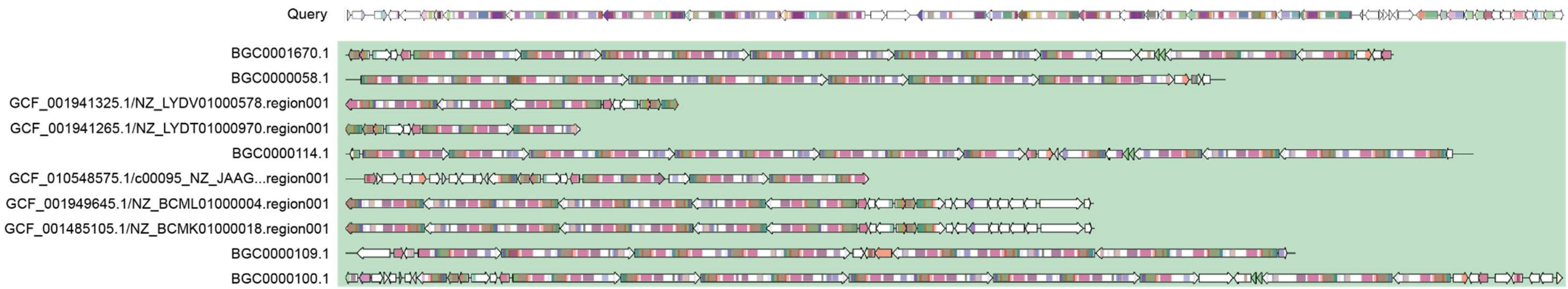


**Fig. 3.** The biosynthetic gene architecture of alligamycin (as Query) from *S. iranensis* and other BGCs from GCF_00315.

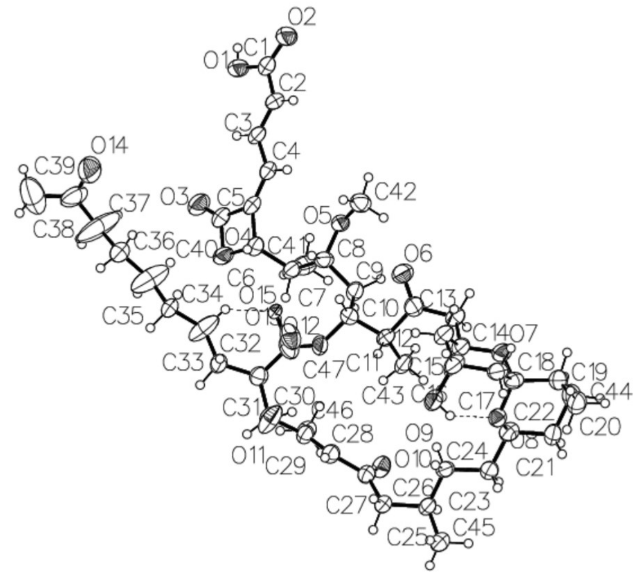


**Fig. 4.** ORTEP drawing of compound **1**.

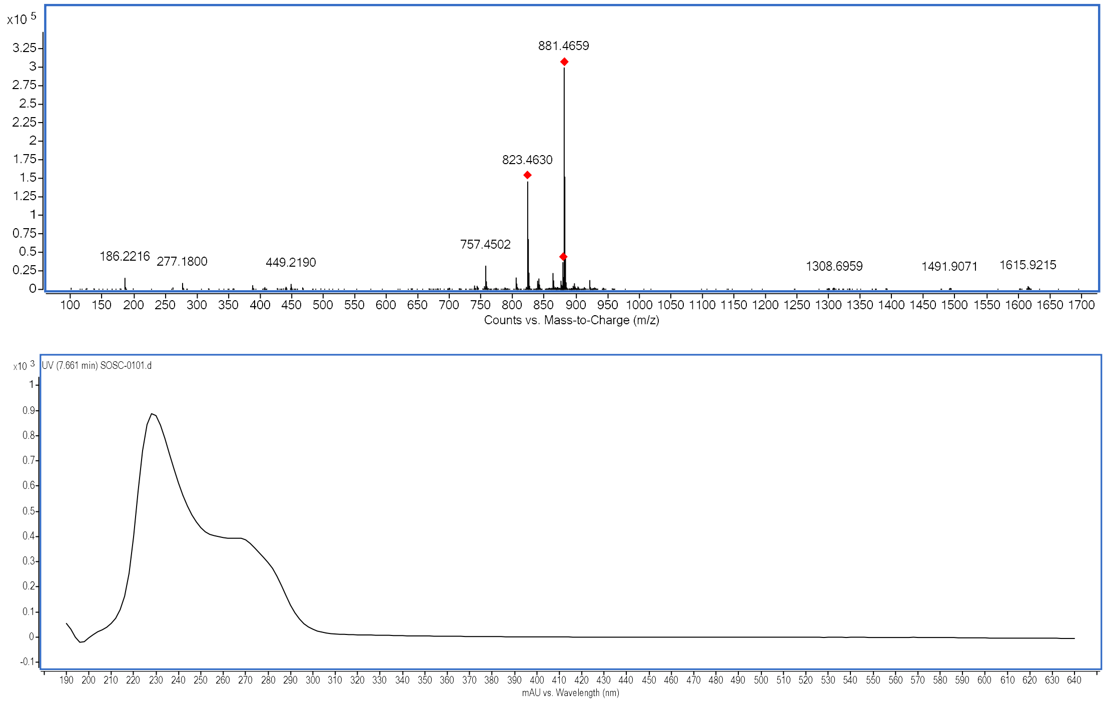


**Fig. 5.** HR-ESI-MS spectrum and UV spectrum of **1**.

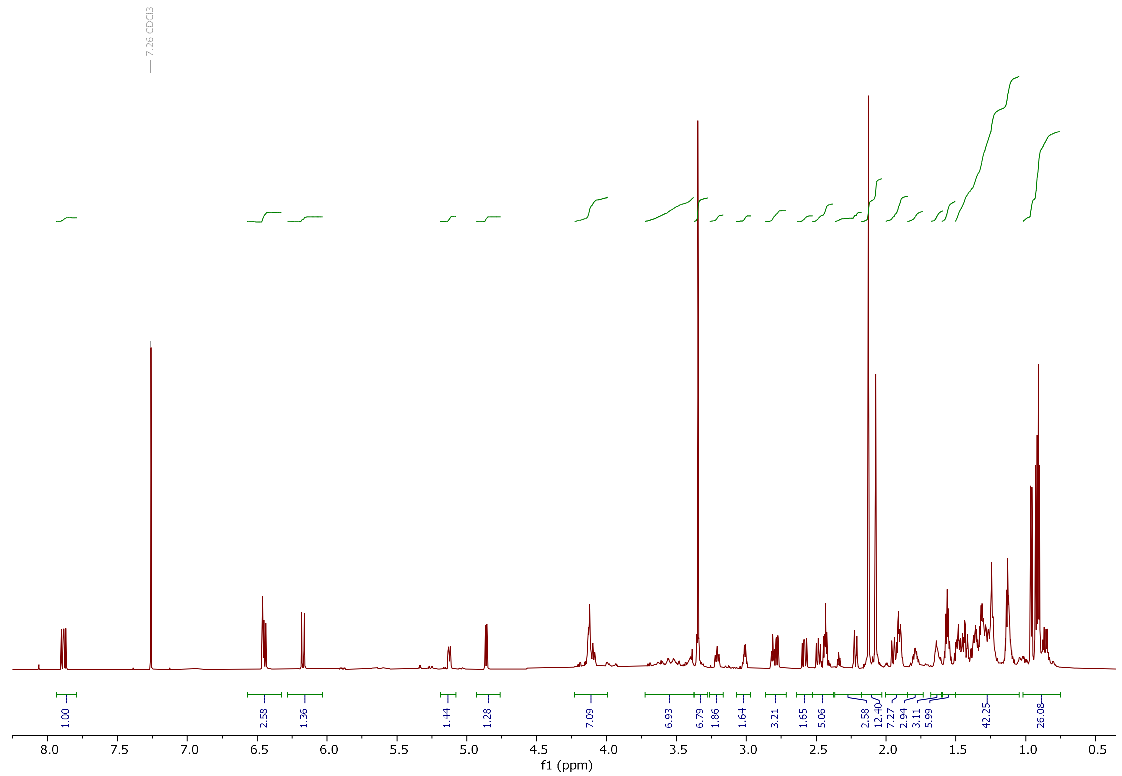


**Fig. 6.** ¹H NMR (800 MHz) spectrum of **1** in $CDCl_3$.

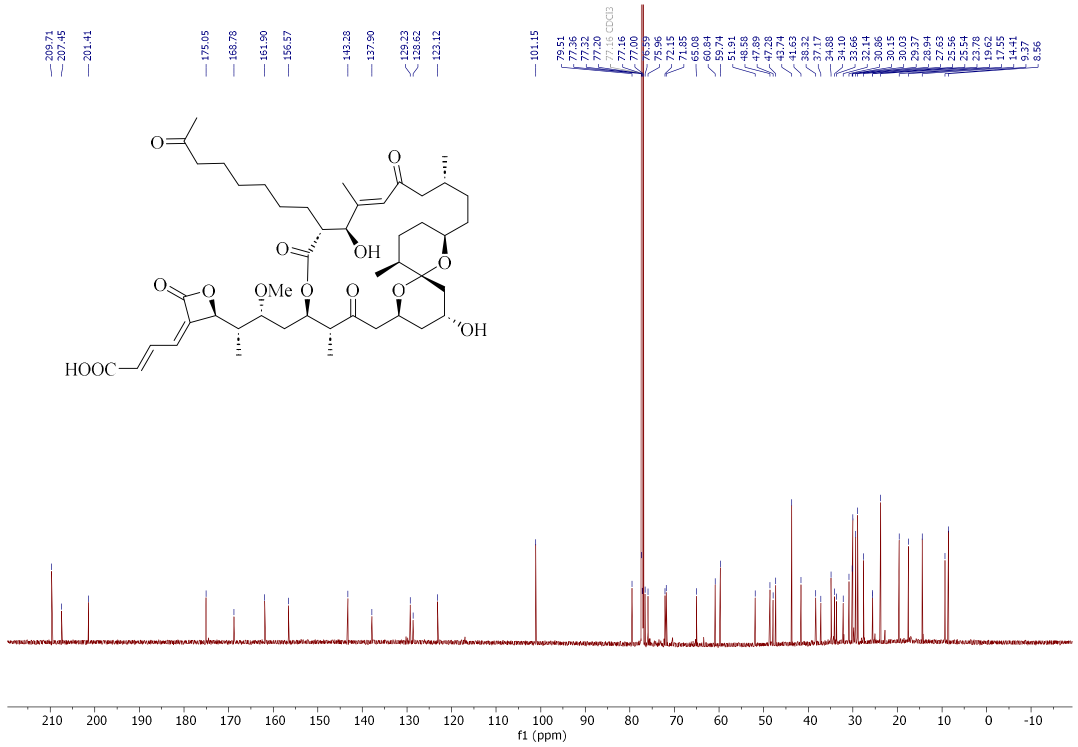


**Fig. 7.** ¹³C NMR (200 MHz) spectrum of **1** in $CDCl_3$.

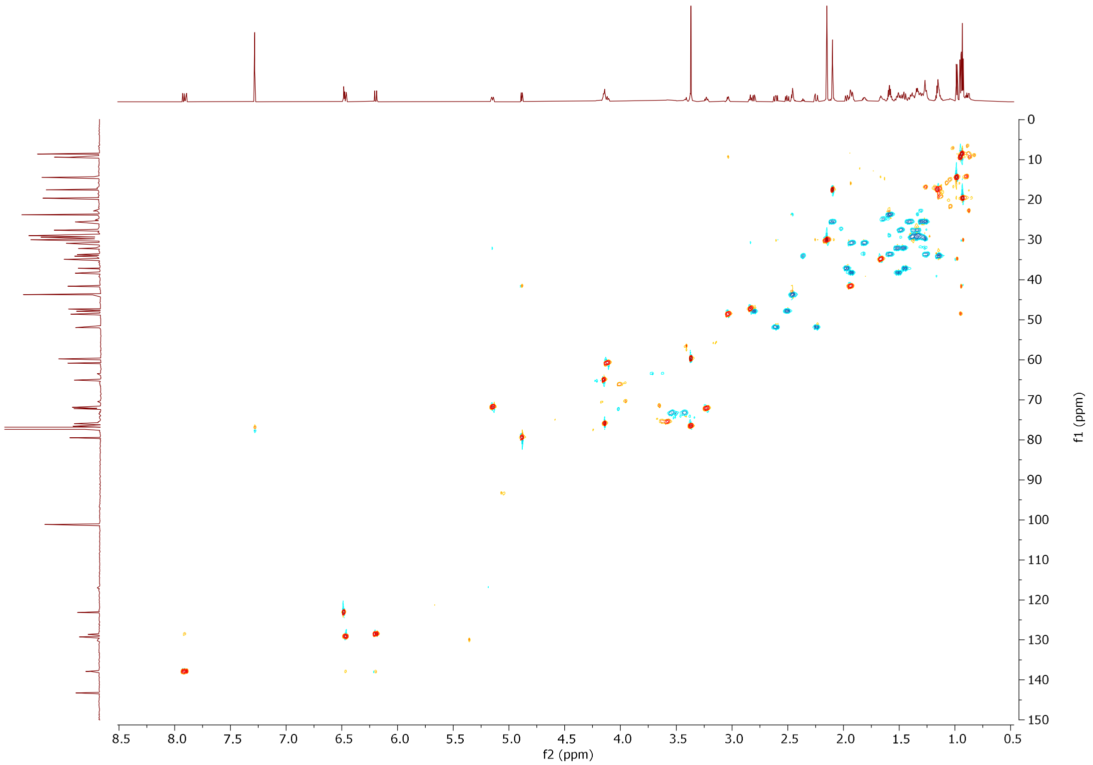


**Fig. 8.** HSQC (800 MHz) spectrum of **1** in $CDCl_3$.

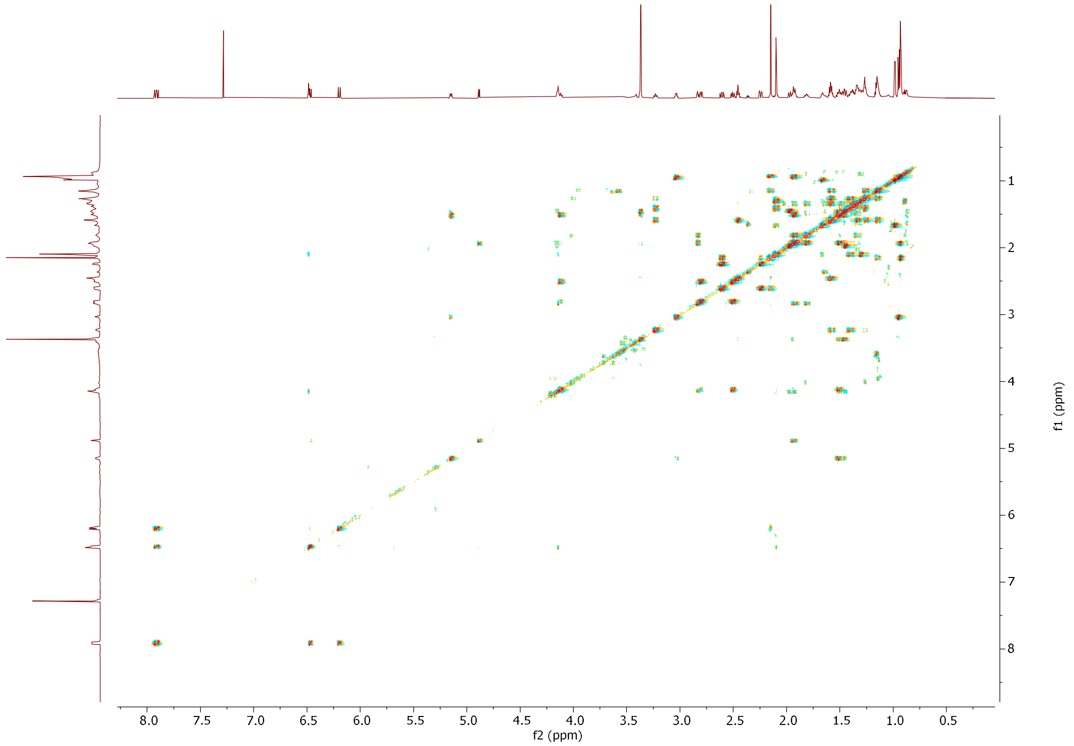


**Fig. 9.** $^{1}H$-$^{1}H$ COSY (800 MHz) spectrum of **1** in $CDCl_3$.

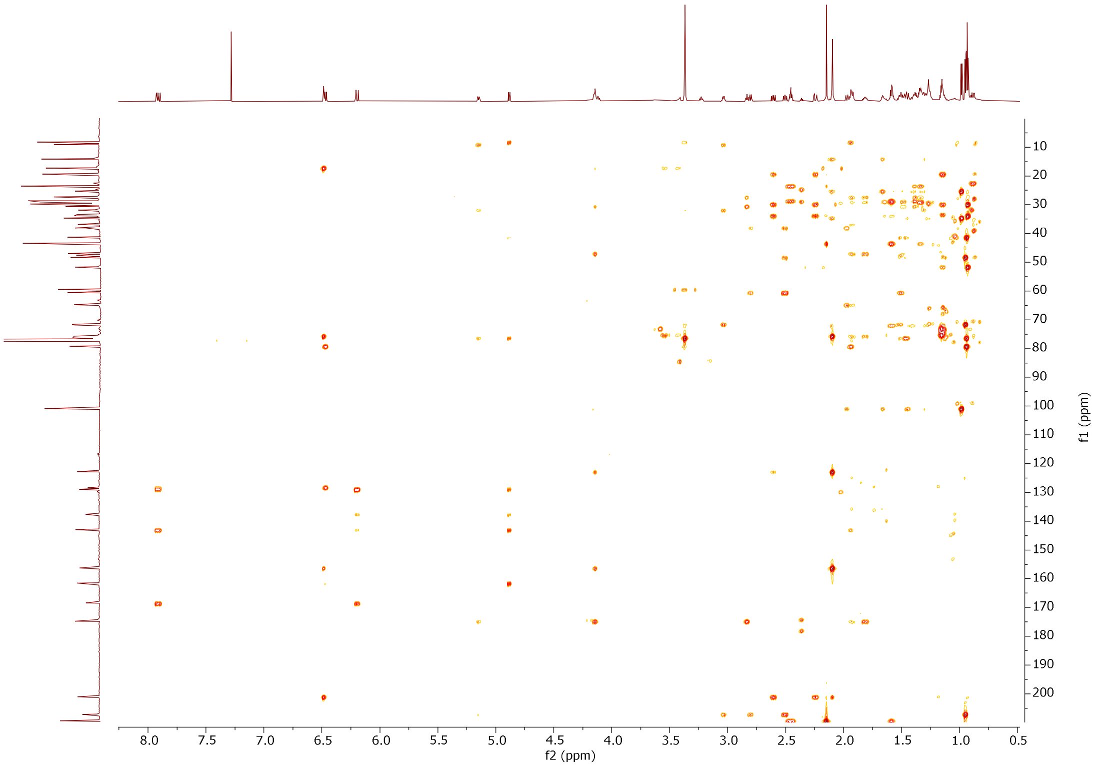


**Fig. 10.** HMBC (800 MHz) spectrum of **1** in $CDCl_3$.

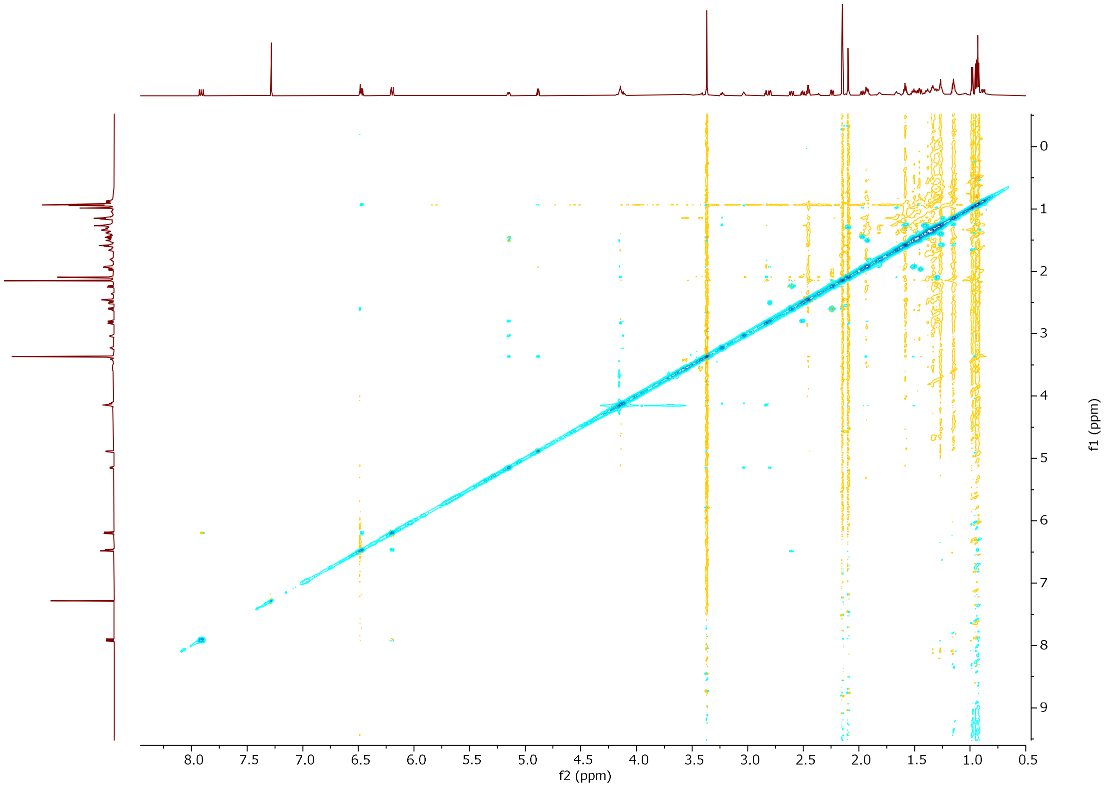


**Fig. 11.** NOESY (800 MHz) spectrum of **1** in $CDCl_3$.

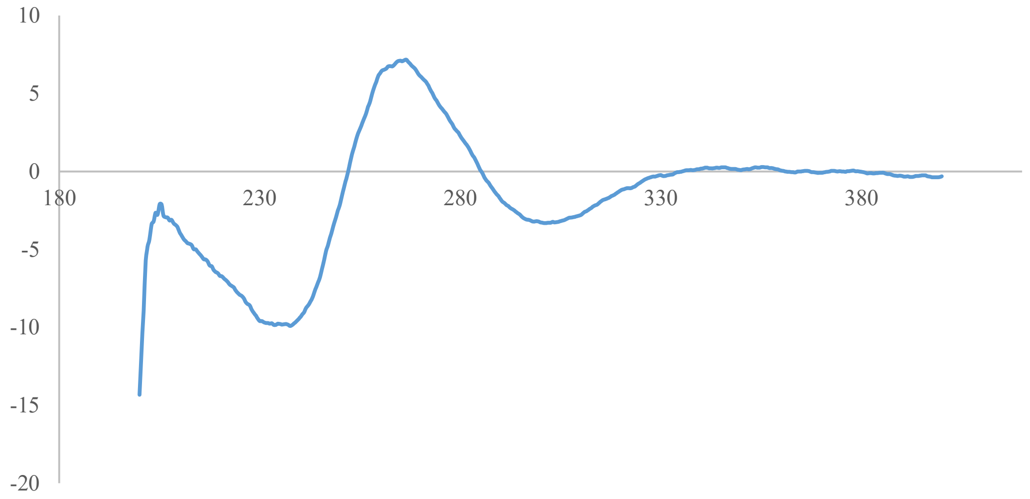


**Fig. 12.** ECD spectrum of **1** (x axis in nm).

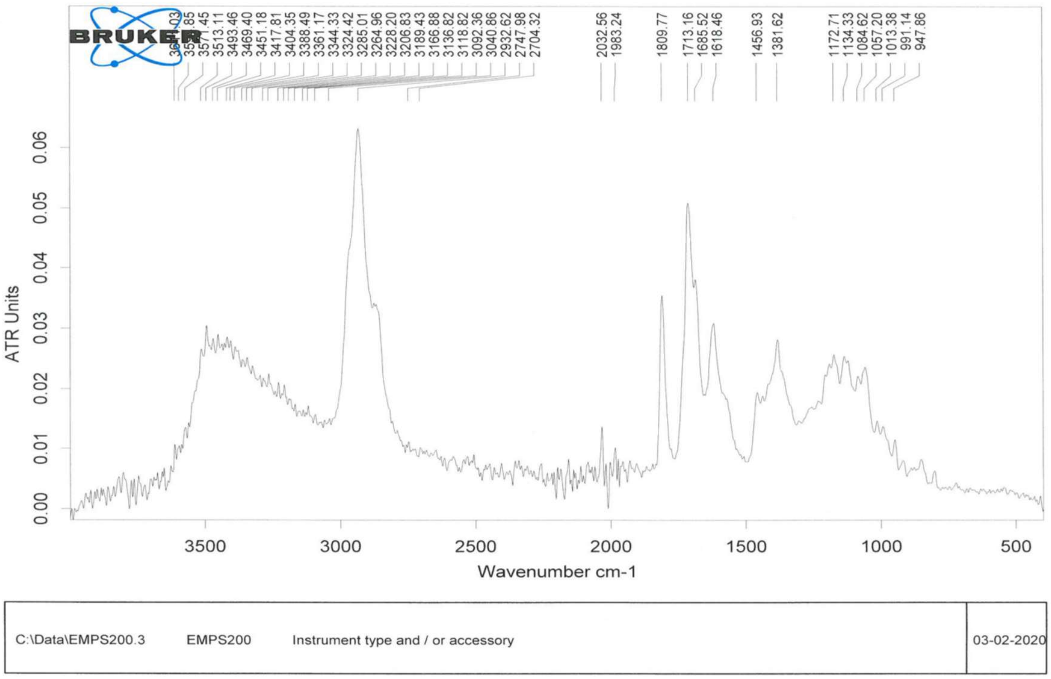


**Fig. 13.** IR spectrum of **1**.

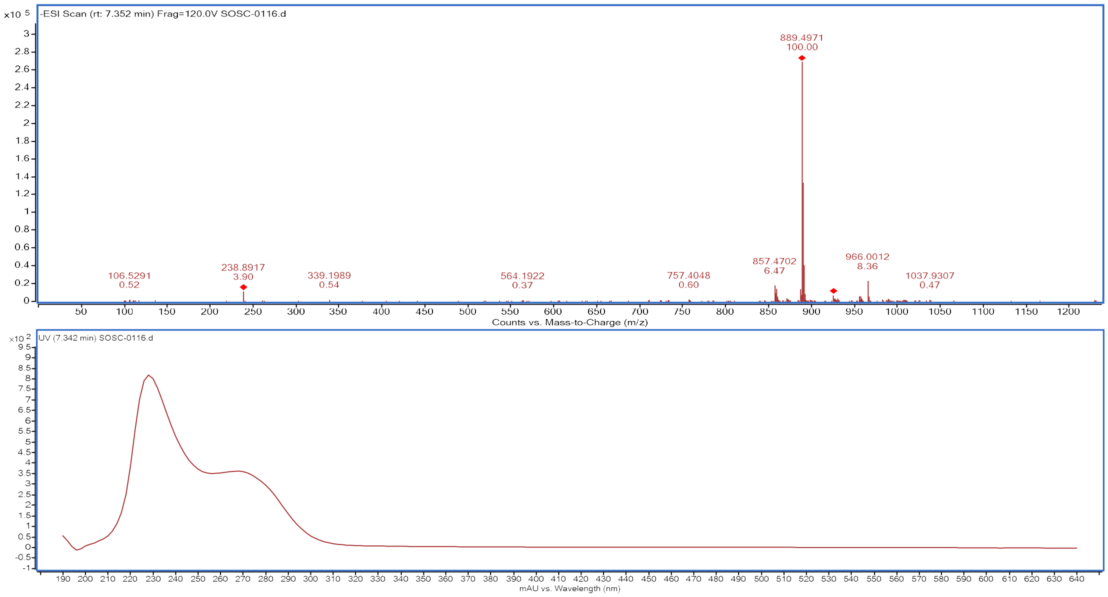


**Fig. 14.** HR-ESI-MS spectrum and UV spectrum of **2**.

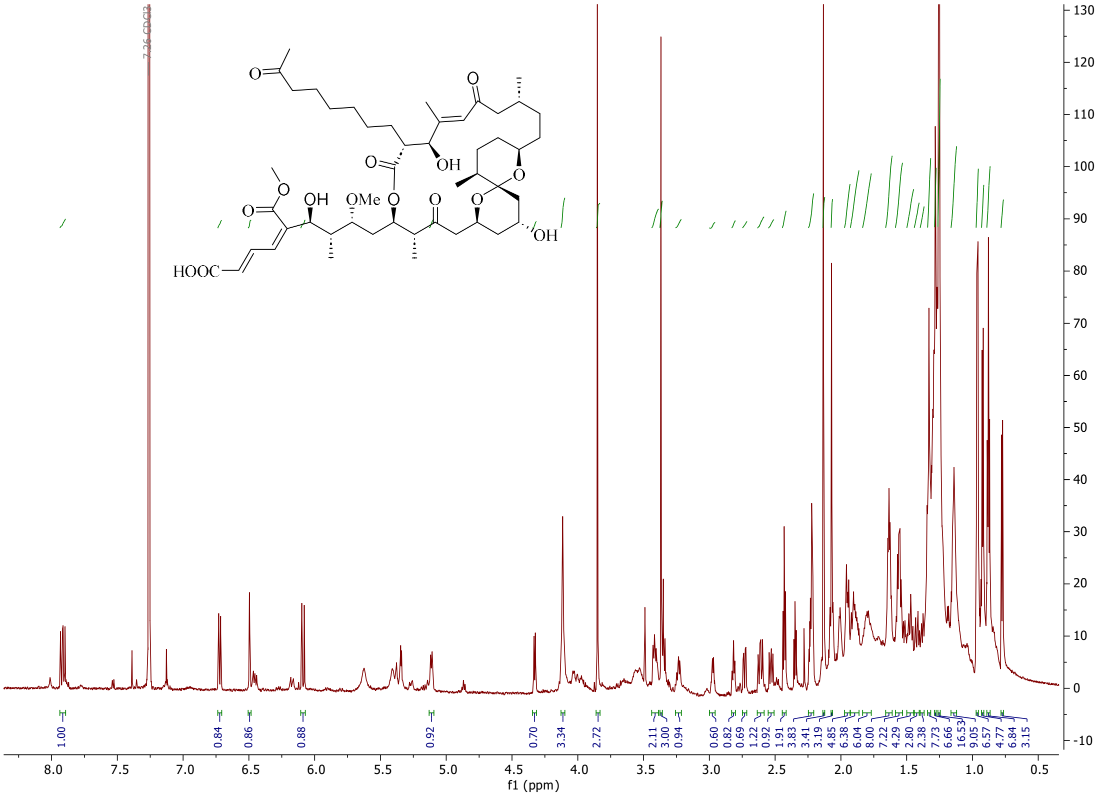


**Fig. 15.** $^{1}$H NMR (800 MHz) spectrum of **2** in $CDCl_3$.

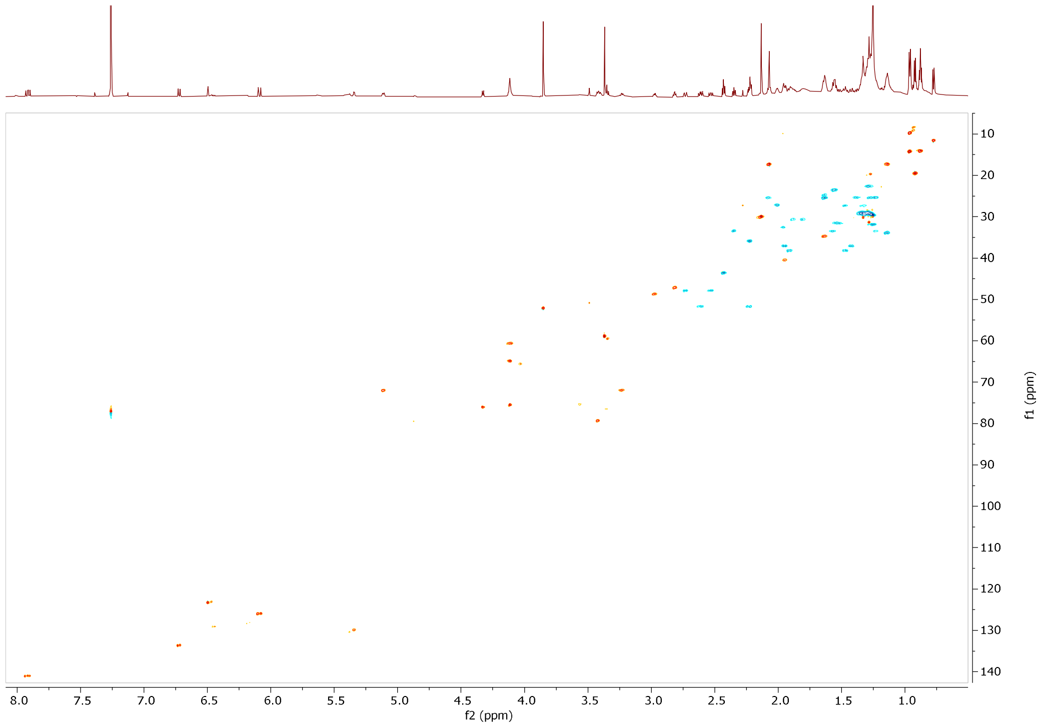


**Fig. 16.** HSQC (800 MHz) spectrum of **2** in $CDCl_3$.

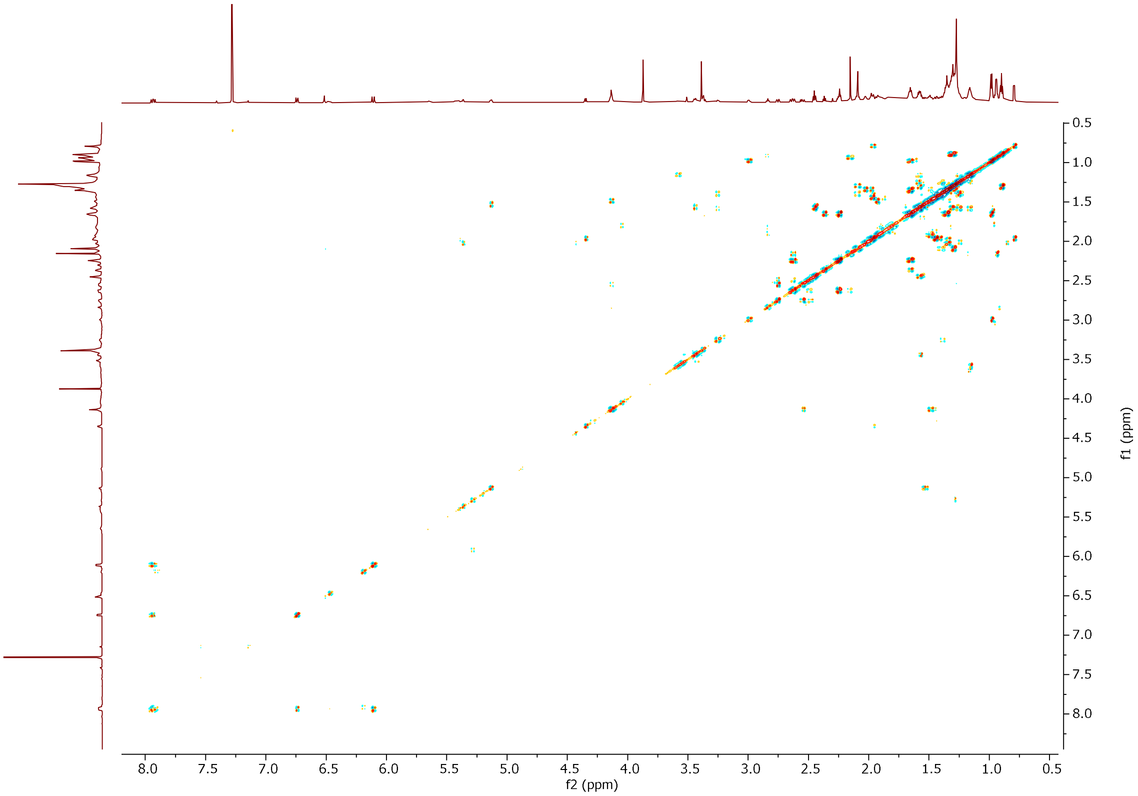


**Fig. 17.** DQF-COSY (800 MHz) spectrum of **2** in $CDCl_3$.

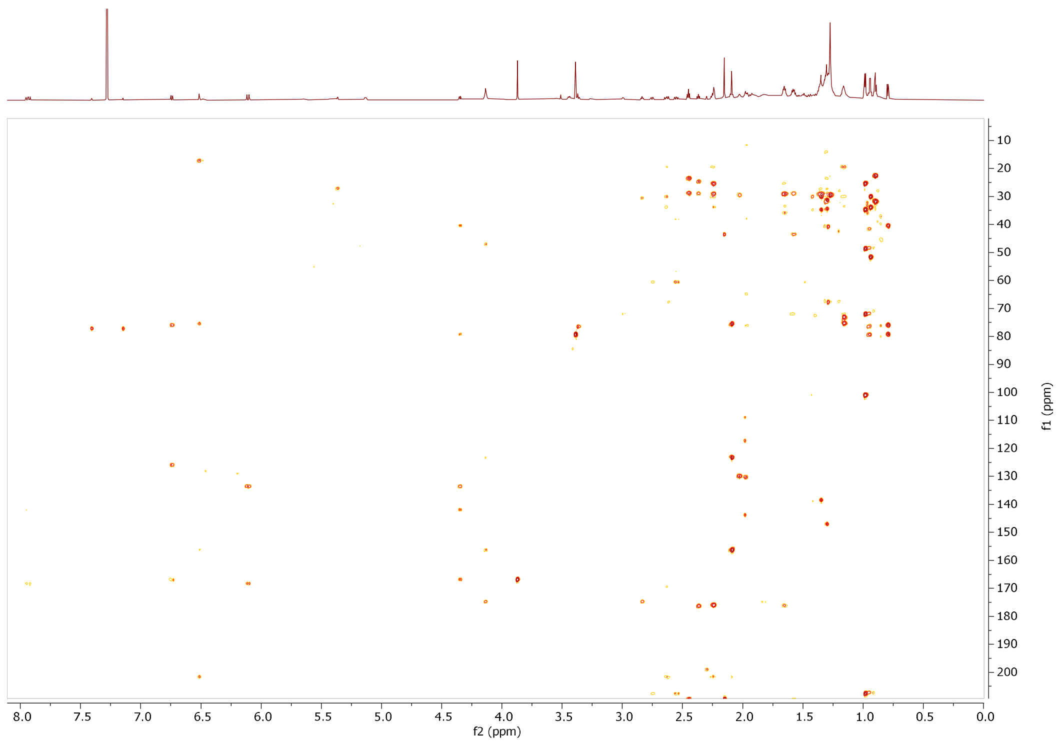


**Fig. 18.** HMBC (800 MHz) spectrum of **2** in CDCl3.

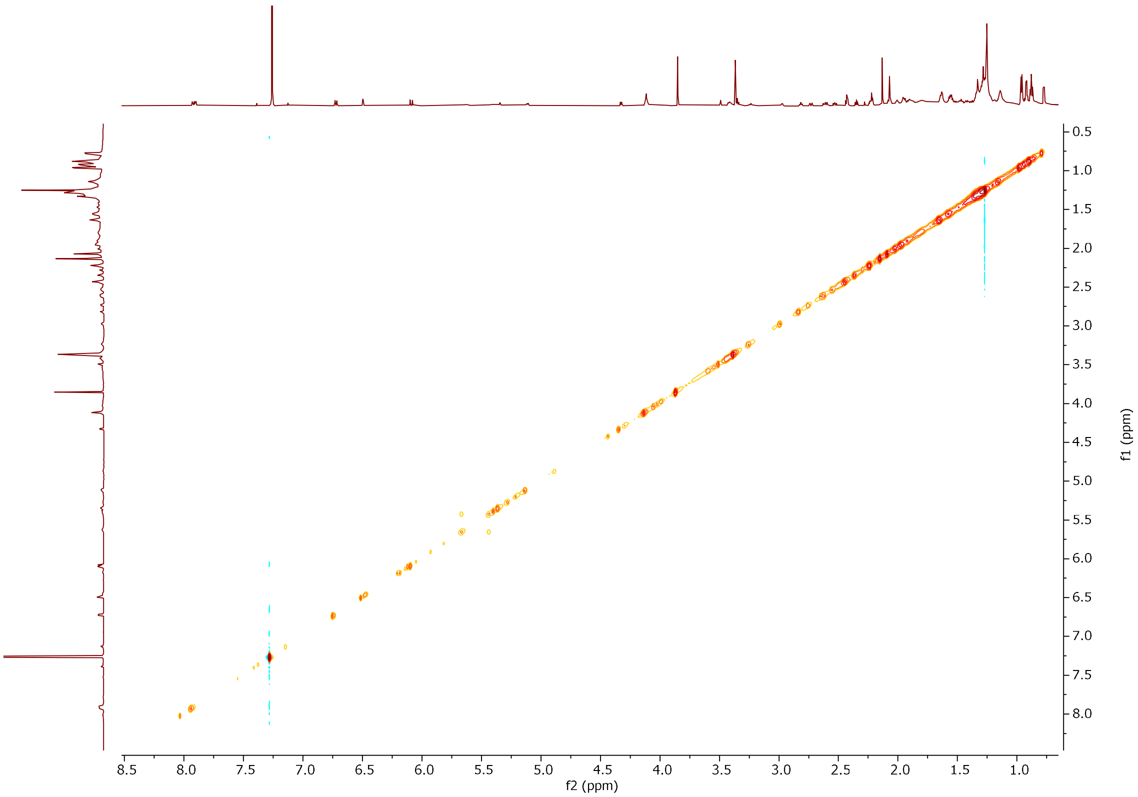


**Fig. 19.** NOESY (800 MHz) spectrum of **2** in CDCl3.

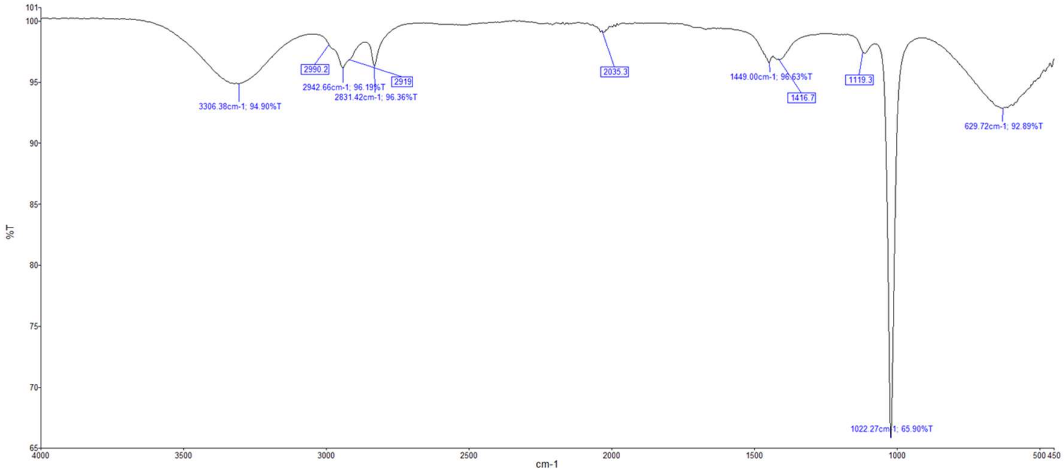


**Fig. 20.** IR spectrum of **2**.

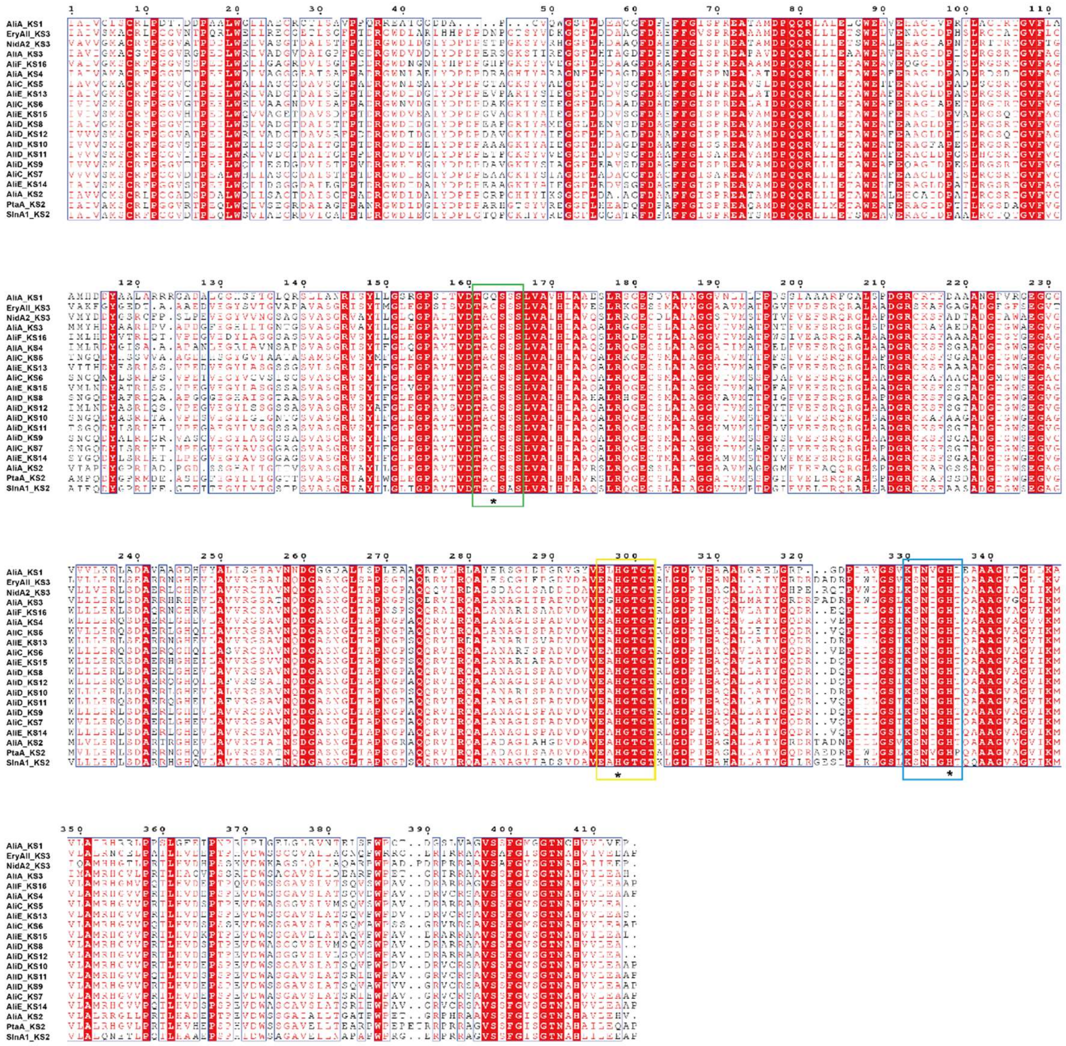

**Fig. 21.** Multiple sequence alignment of KS domains. The catalytic machinery including a cysteine (TACSSS motif in green box) as well as two histidines (EA**H**GTG motif in pink box and KSNIG**H**T motif in blue box) was marked with an asterisk. Abbreviation: Ali, alligamycin; Ery, erythromycin; Nid, niddamycin; Pta, pteridic acid; Sln, salinomycin.

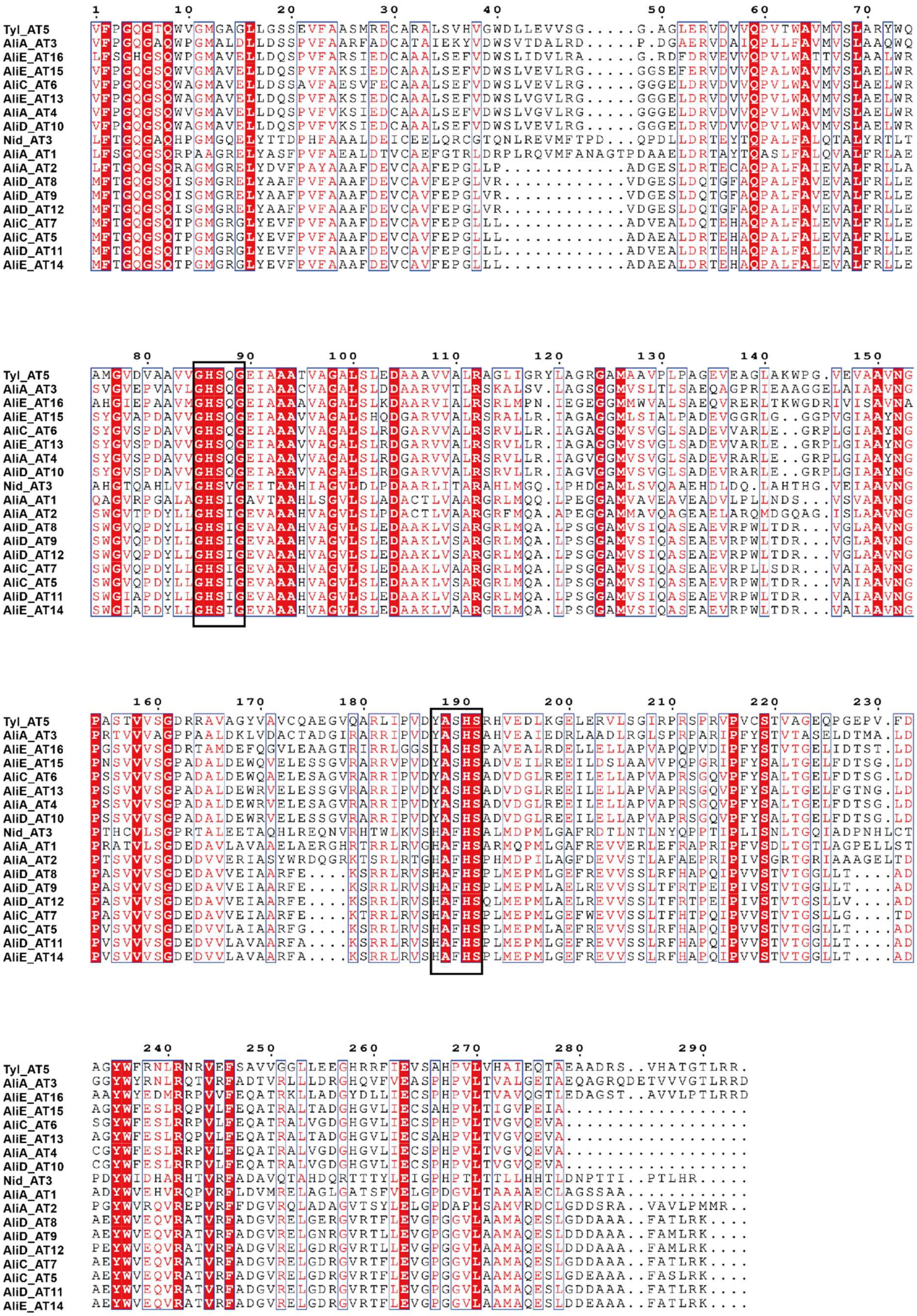

**Fig. 22.** Multiple sequence alignment of AT domains. The black boxes indicate the conserved motifs. Abbreviation: Ali, alligamycin; Nid, niddamycin; Tyl, tylactone.

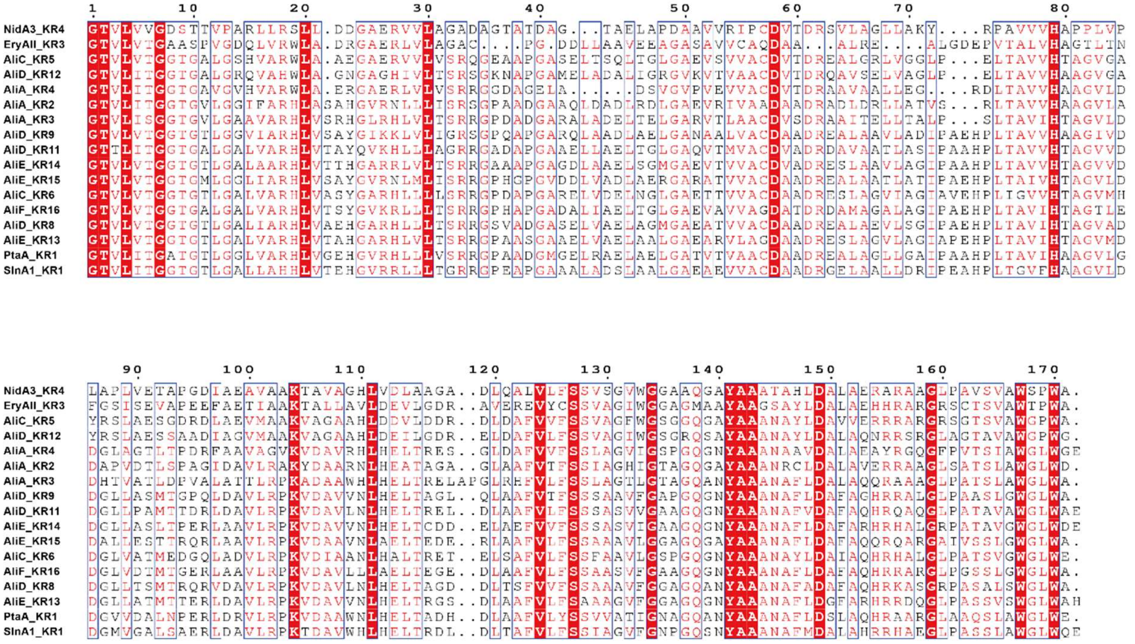


**Fig. 23.** Multiple sequence alignment of KR domains. Abbreviation: Ali, alligamycin; Ery, erythromycin; Nid, niddamycin; Pta, pteridic acid; Sln, salinomycin.

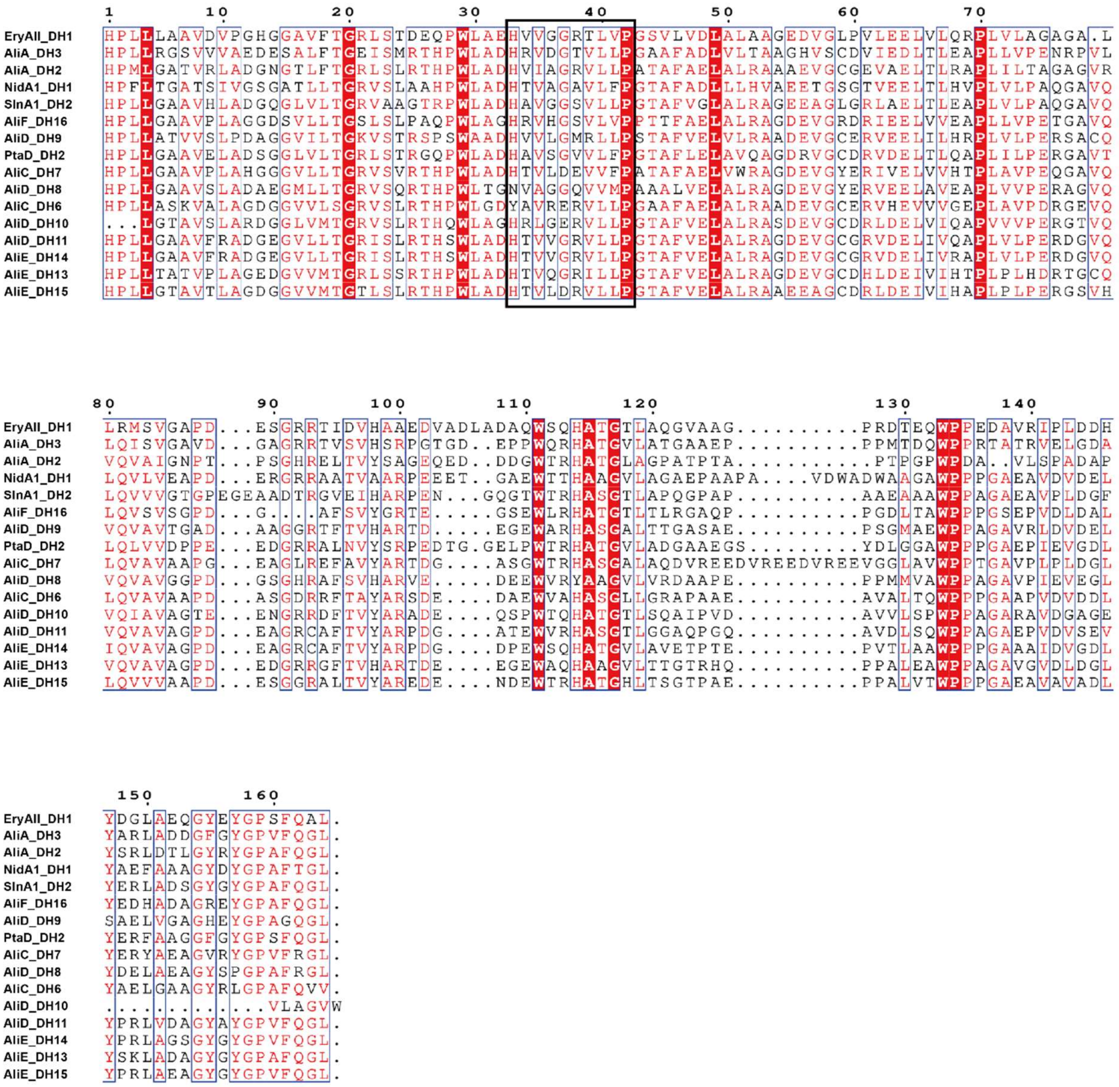

**Fig. 24.** Multiple sequence alignment of DH domains. The black box indicates the conserved HxxxGxxxxP motif. Abbreviation: Ali, alligamycin; Ery, erythromycin; Nid, niddamycin; Pta, pteridic acid; Sln, salinomycin.

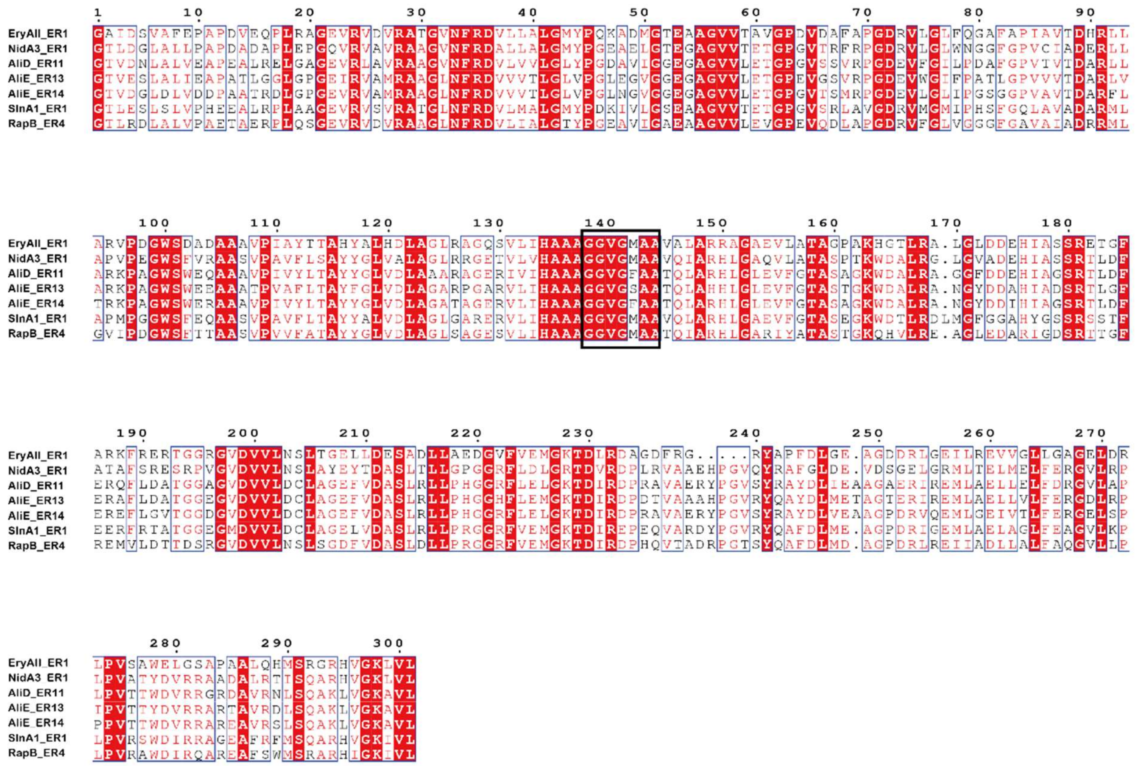

**Fig. 25.** Multiple sequence alignment of ER domains. The unique residue correlated with (2*S*/2*R*)-configuration in the polyketide product is marked with blacked triangles. The position of the NADPH binding site is marked with a black box. Abbreviation: Ali, alligamycin; Ery, erythromycin; Nid, niddamycin; Rap, rapamycin; Sln, salinomycin.

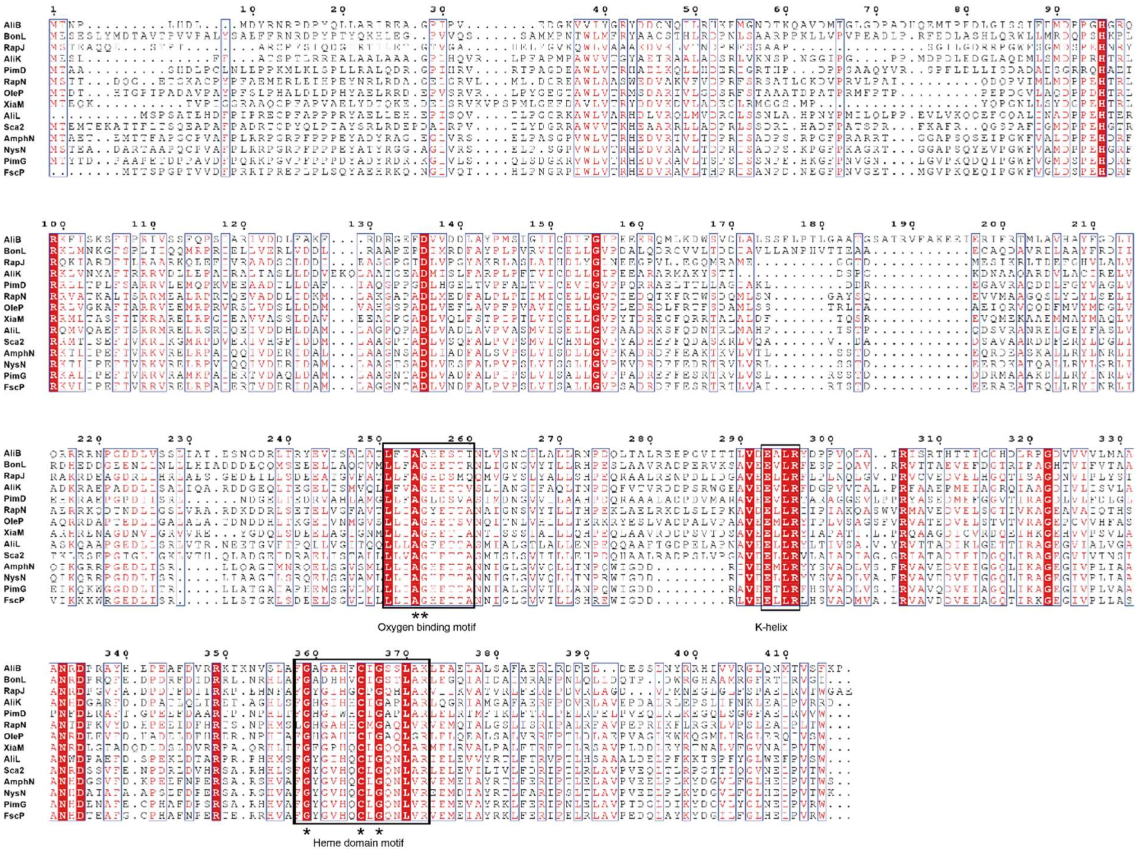


**Fig. 26.** Sequence alignment of AliB, AliK, AliL and other cytochrome P450s. These sequences used in this study: BonL (AFN27475, *Burkholderia gladioli*); SrrO (WP_259329015.1, *Streptomyces rochei*); PimD (CAC20928, *Streptomyces natalensis*); CftA (WP_099966040.1, *Streptomyces* sp. JV178); OleP (AAA92553, *Streptomyces antibioticus*); XiaM (AFK78079, *Streptomyces* sp. SCSIO 02999); Sca2 (BAA06492, *Streptomyces carbophilus*); AmphN (AAK73509, *Streptomyces nodosus*); NysN (AAF71771, *Streptomyces noursei* ATCC 11455); PimG (CAC20928, *Streptomyces natalensis*); FscP (AAQ82557, *Streptomyces* sp. FR-008).

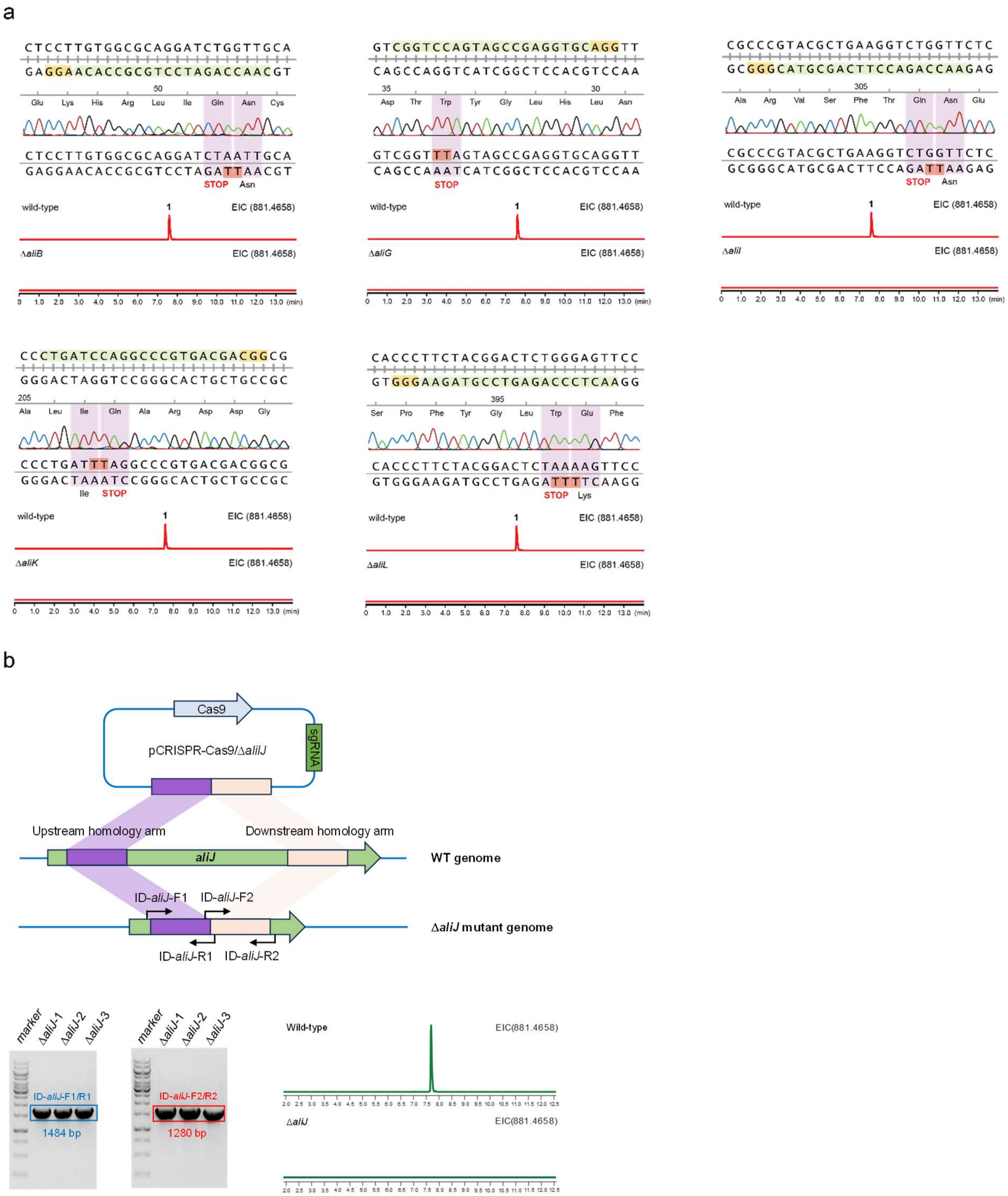


**Fig. 27.** The genome editing of tailoring enzymes in alligamycin biosynthesis. **a**, CRISPR base editing of *aliB*, *aliG*, *aliH*, *aliK*, and *aliL*. **b**, homology directed repair-based CRISRP-Cas9 genome editing.

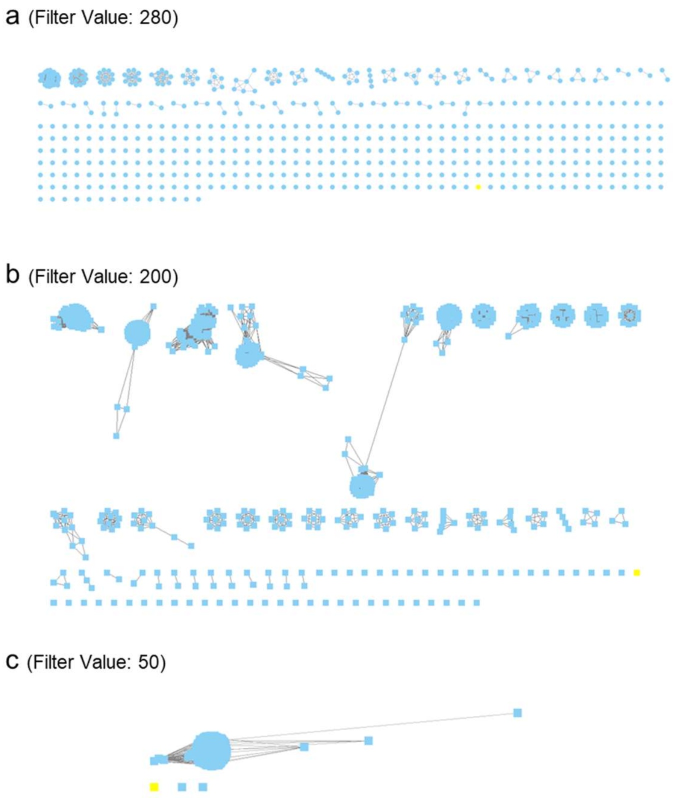


**Fig. 28.** Sequence similarity network of AliJ (colored in yellow). The network was constructed using EFI - enzyme similarity tool (https://efi.igb.illinois.edu/efi-est/).

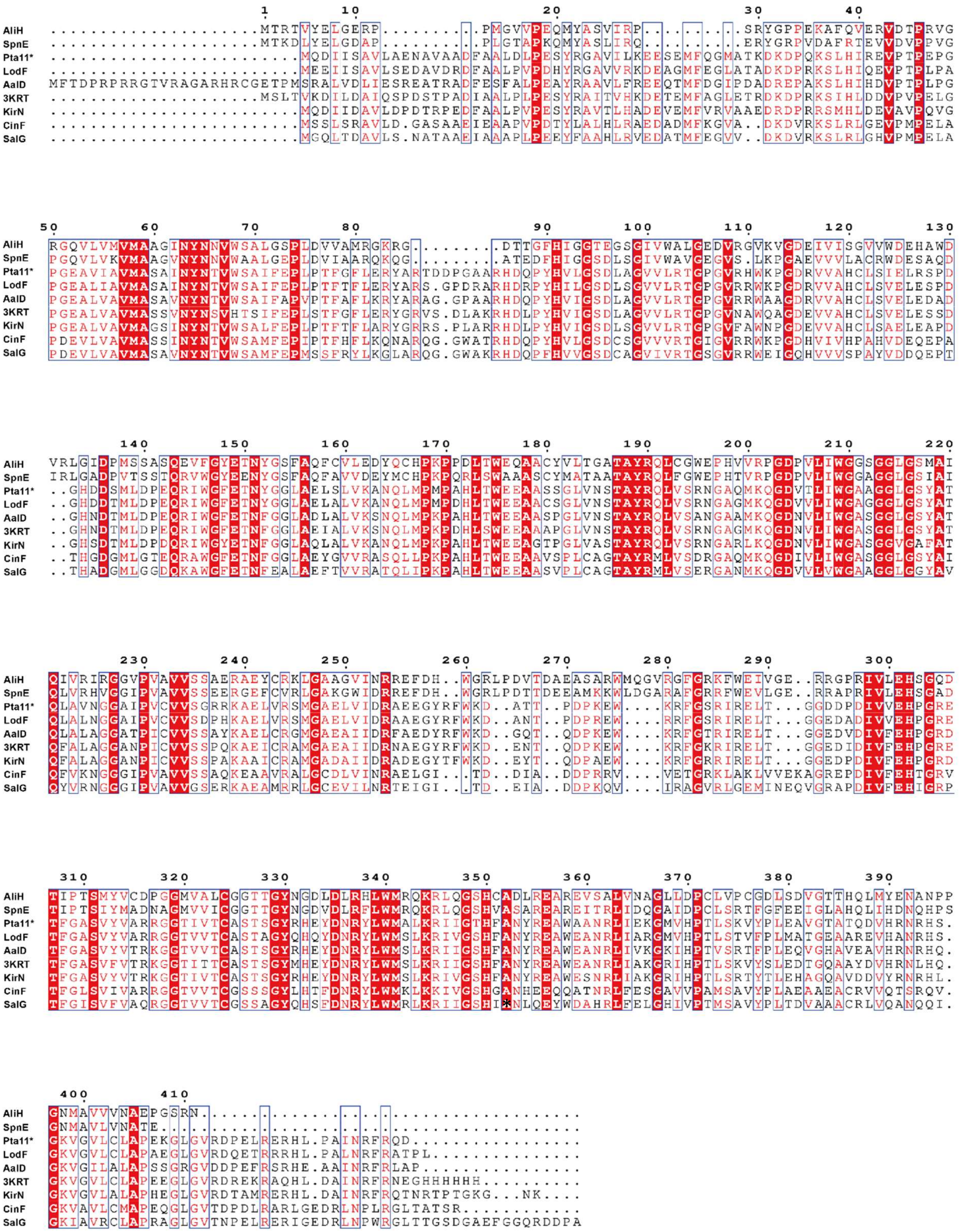

**Fig. 29.** Multiple sequence alignment of AliH and other crotonyl-CoA carboxylase/reductases. These sequences used in this study: SpnE (AKA54628.1, *Streptomyces* sp. CNQ431), Pta11* (WP_044567907.1, *S. iranensis* HM 35), LodF (ANR02547.1, *Streptomyces lasaliensis*), AalD (BBA66510.1, *Streptomyces spiroverticillatus*), 3KRT (*Streptomyces coelicolor*), KirN (CAN89653.1, *Streptomyces collinus* Tu 365), CinF (CBW54676.1, *Streptomyces cinnabarigriseus*), SalG (ABP73651.1, *Salinispora pacifica*).

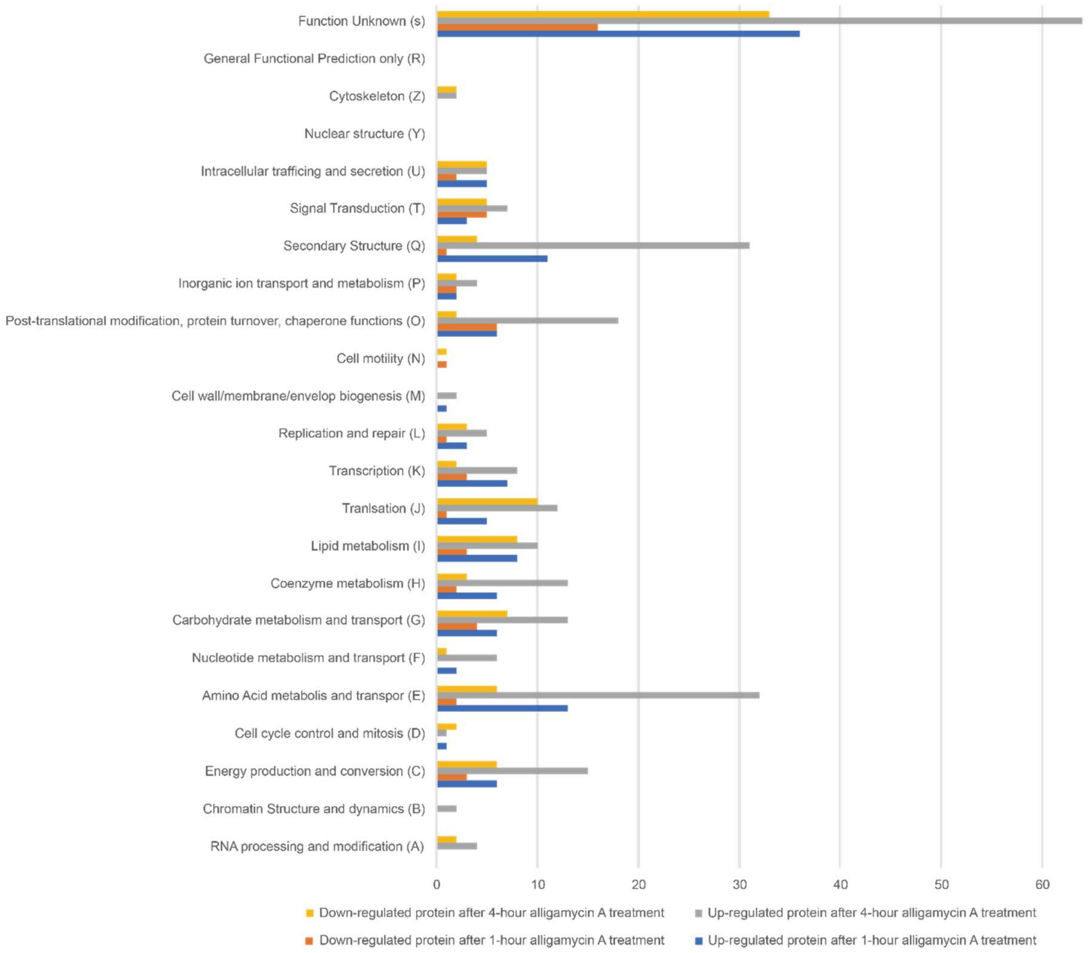


**Fig. 30.** The COGs functional classification of differential proteins in *A. niger*.

a

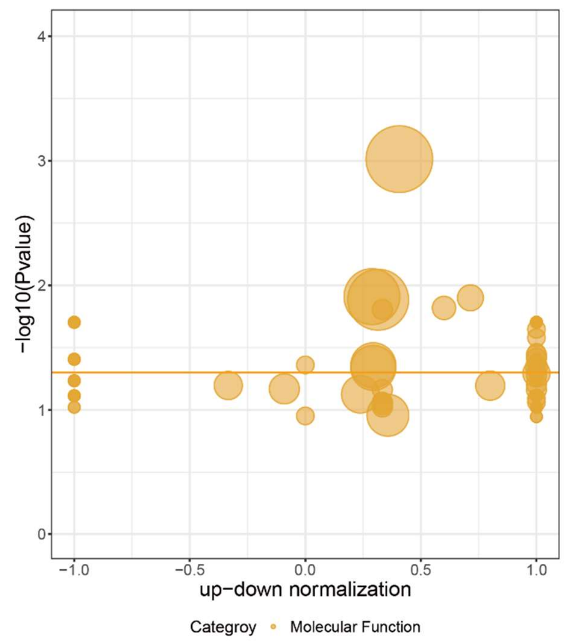


GO term Top 20

| ID | Descrption |
|---|---|
| GO:0003824 | catalytic activity |
| GO:0043167 | ion binding |
| GO:0016874 | ligase activity |
| GO:0005488 | binding |
| GO:0019842 | vitamin binding |
| GO:0004252 | serine−type endopeptidase activity |
| GO:0003863 | 3−methyl−2−oxobutanoate dehydrogenase (2−methylpropanoyl−transferring) activity |
| GO:0003972 | RNA ligase (ATP) activity |
| GO:0004037 | allantoicase activity |
| GO:0004339 | glucan 1,4−alpha−glucosidase activity |
| GO:0004694 | eukaryotic translation initiation factor 2alpha kinase activity |
| GO:0004788 | thiamine diphosphokinase activity |
| GO:0004851 | uroporphyrin−III C−methyltransferase activity |
| GO:0008452 | RNA ligase activity |
| GO:0008531 | riboflavin kinase activity |
| GO:0018506 | maleylacetate reductase activity |
| GO:0019789 | SUMO transferase activity |
| GO:0030975 | thiamine binding |
| GO:0051730 | GTP−dependent polyribonucleotide 5'−hydroxyl−kinase activity |
| GO:0051732 | polyribonucleotide kinase activity |

b

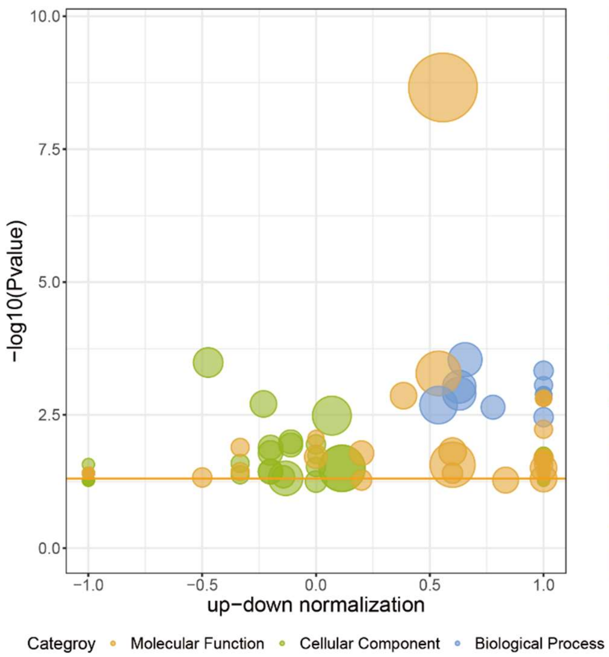


GO term Top 20

| ID | Descrption |
|---|---|
| GO:0003824 | catalytic activity |
| GO:0005739 | mitochondrion |
| GO:0006082 | organic acid metabolic process |
| GO:0006144 | purine nucleobase metabolic process |
| GO:0072523 | purine−containing compound catabolic process |
| GO:0019752 | carboxylic acid metabolic process |
| GO:0043436 | oxoacid metabolic process |
| GO:0009448 | gamma−aminobutyric acid metabolic process |
| GO:0019628 | urate catabolic process |
| GO:0046415 | urate metabolic process |
| GO:0016491 | oxidoreductase activity |
| GO:0016874 | ligase activity |
| GO:0004053 | arginase activity |
| GO:0004846 | urate oxidase activity |
| GO:0016663 | oxidoreductase activity, acting on other nitrogenous compounds as donors, oxygen as acceptor |
| GO:0044281 | small molecule metabolic process |
| GO:1901565 | organonitrogen compound catabolic process |
| GO:0044429 | mitochondrial part |
| GO:0005737 | cytoplasm |
| GO:0009112 | nucleobase metabolic process |

**Fig. 31.** The GO annotation of differential proteins in *A. niger*. ***a***, 1-hour treatment with alligamycin A. ***b***, 4-hour treatment of alligamycin A.

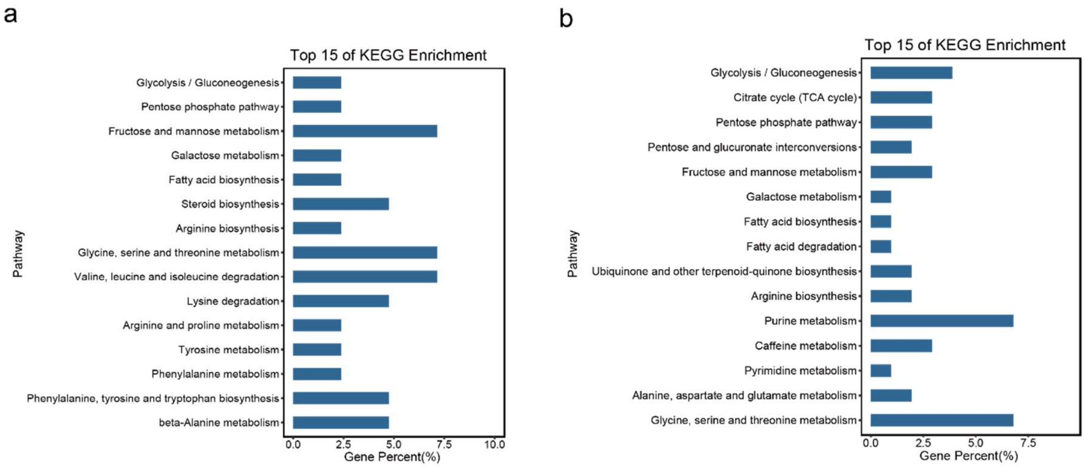


**Fig. 32.** The KEGG pathway enrichment analysis of differential proteins in *A. niger*. ***a***, 1-hour treatment with alligamycin A. ***b***, 4-hour treatment of alligamycin A.

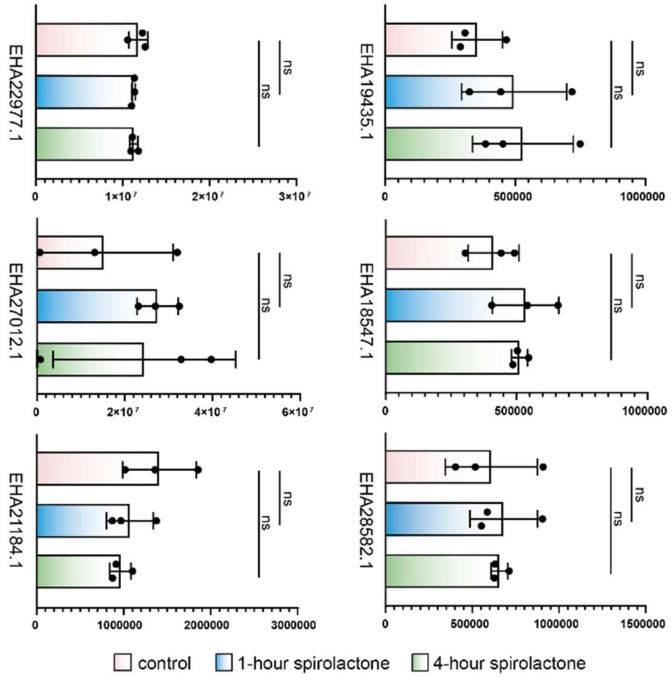


**Fig. 33.** The abundance of responsive protein of known antifungal drugs in alligamycin A treatment. EHA22977.1, Erg13; EHA27012.1, manganese-superoxide dismutase; EHA21184.1, squalene epoxidase; EHA19435.1, CYP51; EHA18547.1, 1,3-β-D-glucan synthase (EHA18547.1); EHA28582.1, ChiA1-like chitinase. Statistical significance was assessed by Student's t-test. Abbreviation: *ns*, not significant.
